# Context-dependent response of endothelial cells to *PIK3CA* mutation

**DOI:** 10.1101/2025.02.25.640041

**Authors:** Helena Sabata, Ariadna Roca-Coll, Jose A. Dengra, Leonor Gouveia, Ane Martinez-Larrinaga, Alberto Collado-Remacha, Sandra D. Castillo, Elena Castillo, Nathalie Tisch, Macarena De Andrés-Laguillo, Svanhild Nornes, Judith Llena, Pilar Villacampa, Bart Vanhaesebroeck, María P. Alcolea, Sarah De Val, Susana Lopez-Fernandez, Katrien De Bock, Eulalia Baselga, Rui Benedito, Ana Angulo-Urarte, Mariona Graupera

## Abstract

Cancer mutations in the *PIK3CA* gene cause congenital disorders. The endothelium is among the most frequently affected tissues in these disorders, displaying aberrant vascular overgrowth in the form of malformations. Pathological *PIK3CA* vascular phenotypes are found in veins and capillaries but rarely in arteries for reasons that are unclear at present. Here, using lineage tracing, we show that expression of mutated *PIK3CA^H1047R^* in endothelial cells leads to marked clonal expansions in capillary and venous endothelial cells. In contrast, mature arterial endothelial cells are refractory to *PIK3CA* mutation under these conditions and never display pathological phenotypes. Moreover, *PIK3CA^H1047R^*expression in arterial precursors interrupts arterial differentiation, thereby driving fate switch towards venous identity. This fate rewiring offers an additional layer of protection to prevent arterial damage in response to *PIK3CA* genetic perturbation. Molecularly, the *PIK3CA^H1047R^*-driven arterial-to-venous fate switch is orchestrated by upregulation of the vein-specifying transcription factor *Nr2f2*/COUP-TFII. Our findings reveal that pathogenic responses to *PIK3CA^H1047R^*greatly depend on the diferentation stage and fate trajectory of the targeted cell. Arteries are thus shielded against *PIK3CA* mutation, solving the long-standing question on the rarity of *PIK3CA*-related arterial malformations observed in patients.

## Introduction

Somatic and germline pathogenic variants in genes associated with cancer have been identified as a cause of a variety of congenital disorders (Castel, Rauen, and McCormick 2020; Angulo-Urarte and Graupera 2022; Nussinov, Tsai, and Jang 2022; Samuels et al. 2004). In these conditions, mutations in the germline can be either inherited or appear *de novo*. Instead, somatic mutations arise during embryonic development, resulting in mosaic distribution and a broad spectrum of phenotypes (Castel, Rauen, and McCormick 2020). Recent findings have shown that cancer mutations also exist in phenotypically normal tissues (Herms and Jones 2023; Martincorena et al. 2018). Understanding why and how these mutations result in context-dependent manifestations is crucial for identifying tissue susceptibility to pathogenic outcomes and explaining the diversity of phenotypes observed in patients.

*PIK3CA* encodes the p110α phosphoinositide 3-kinase (PI3K) catalytic subunit (hereafter referred to PI3Kα) that regulates cell growth, proliferation, migration, and metabolism (Vanhaesebroeck et al. 2010), mainly through activation of AKT-mTOR signaling (Hoxhaj and Manning 2020; Manning and Toker 2017). *PIK3CA* pertains to the group of oncogenes frequently mutated in a mosaic fashion in congenital disorders and normal tissues. Activating mutations in *PIK3CA* span the entire gene, albeit there are common hotspots across conditions (E545K, E542K, H1047R, H1047L) (Angulo-Urarte and Graupera 2022). Mesodermal derivatives such as the endothelium are among the tissues most affected in *PIK3CA*-related congenital disorders, manifesting in aberrant overgrowth and malformations (Angulo-Urarte and Graupera 2022; Madsen, Vanhaesebroeck, and Semple 2018). While tissues carrying *PIK3CA* mutations in the epithelium have been extensively studied (Van Keymeulen et al. 2015; Koren et al. 2015; Herms et al. 2024; Yum et al. 2021), the pathogenic responses of the mesodermal lineage in general, and endothelial cells in particular, to these mutations are much less known.

Vascular malformations are abnormal vascular channels that may occur anywhere in the body and strongly impact the patient’s quality of life. These vascular lesions appear during embryonic development and continue growing proportionally throughout the patient’s lifetime. They are classified as slow-flow lesions if they appear in capillaries, veins, and lymphatic vessels and fast-flow lesions when malformations affect arteries (International Society for the Study of Vascular Anomalies 2018). *PIK3CA* mutations have been identified in the endothelium of all types of slow-flow vascular malformations (*i.e.,* capillary, venous, and lymphatic malformations). These lesions form through increased cell proliferation, although *PIK3CA* mutations alone do not confer a proliferative advantage to endothelial cells (ECs). Expansion of the mutant cells selectively occurs in the presence of mitogenic signals, which explains why *PIK3CA* mutations in ECs exhibit detrimental effects during organ growth and hormonal spurs (Kobialka et al. 2022; Hassanein et al. 2012; Petkova et al. 2023; Martinez-Corral et al. 2020). Currently, there is no evidence that *PIK3CA* mutations cause fast-flow vascular malformations (Angulo-Urarte and Graupera 2022). Thus far, it remains unknown whether arterial cells harbor *PIK3CA* mutations and are resistant to them or whether the lack of pathological traits in these vessels is because mutant clones are negatively selected or due to alterations in differentiation pathways.

Retinal blood vessels in mice emerge postnatally in a process known as sprouting angiogenesis (Selvam, Kumar, and Fruttiger 2018). During the first week of the angiogenesis, *de novo* vessels grow in a 2D fashion from the optic nerve and require the secretion of mitogenic endothelial factors. The sprouting process involves the selection of two molecularly distinct cell types: tip cells, which function as leaders of the sprout and primarily migrate, and stalk cells, which facilitate sprout elongation through proliferation (Blanco and Gerhardt 2013; Potente, Gerhardt, and Carmeliet 2011). A subset of tip cells expresses arterial markers, migrates backward against the flow, and contributes to the growth of developing arteries (Xu et al. 2014; Pitulescu et al. 2017). Retinal vessels are amenable to whole-mount analysis, which allows the tracing of cell behaviors in specific EC subsets (*i.e.,* tip, arterial, capillary, vein) at single-cell resolution. Based on these unique features, retinal vessels represent an ideal system to interrogate why the same mutation results in distinct phenotypic outcomes in different EC subsets.

Here, to gain insight into why pathogenic phenotypes are restricted to some EC types, we map the behavior of cells bearing *PIK3CA* mutations at clonal resolution. We use a compendium of genetic approaches that allow the faithful tracing of *PIK3CA^H1047R^*mutant clones in different EC subsets. We further use single-cell RNA-sequencing analysis to characterize how *PIK3CA^H1047R^* impacts endothelial cell fate trajectories leading to context-dependent pathological phenotypes in the vasculature.

## Results

### Spatial mapping of *Pik3ca^H1047R^*-driven clonal expansion in the retinal vasculature

Expression of the *Pik3ca^H1047R^* allele (*Pik3ca^tm1.1Waph/+^)* in mouse retinal ECs using the *Pdgfb-CreER^T2^* mouse line results in venous and capillary malformations while arteries remain unperturbed (**Extended Data Fig. 1a-c**) (Kobialka et al. 2022). To trace how these phenotypes correlate with the presence or absence of mutant cells, we crossed the *R26-mTmG* reporter allele into the *Pik3ca^H1047R^*; *Pdgfb-CreER^T2^* mice (**Fig. 1a,b**). In these mice, tamoxifen treatment stochastically activates Cre in any EC subset, followed by co-expression of the *Pik3ca^H1047R^* allele from its endogenous promoter and a membrane-bound fluorescent GFP reporter that provides an approximation of mutant *Pik3ca* expressing cells. *Pdgfb-CreER^T2^; R26-mTmG* mice were used as controls (hereafter referred to as *pEC-GFP)*. We treated pups at postnatal day (P) 1 with a low dose of 4-hydroxytamoxifen (4-OHT) and isolated retinas at P6 (**Fig. 1c**). We obtained a mosaic recombination pattern in *pEC-GFP* and *pEC-GFP-Pik3ca^H1047R^*retinas with GFP+ cells scattered throughout the vascular plexus (**Fig. 1d**). First, we analyzed the incidence of vascular malformations in *pEC-GFP-Pik3ca^H1047R^* retinas by identifying aberrant overgrown vessels composed of *Pik3ca^H1047R^*-GFP+ ECs. We observed a very high incidence of malformed vessels in capillaries and veins (87% and 62%, respectively) but none in arteries (**Fig. 1e**). High magnification images confirmed that wild-type and mutant GFP+ ECs were present in all vessel types (**Fig. 1f**) but that *Pik3ca^H1047R^*-GFP+ ECs only expanded in capillaries and veins (**Fig. 1f,g**). Expansion of mutant clones led to aberrant vascular tubes with increased vessel width (**Fig. 1f,h**). We confirmed that this expansion was associated with increased EC proliferation at the onset of the malformation (**Extended Data Fig. 1d-f**). Despite *Pik3ca^H1047R^*-GFP+ cells in arteries, these mutant cells did not expand or alter arterial vessel width (**Fig. 1f-h**). We monitored PI3K/AKT/mTORC1 signaling activation by staining for phospho (p)-S6 (Ser235/236) and found that capillary and venous *Pik3ca^H1047R^*-GFP+ ECs exhibited higher activation of PI3K signaling compared to their control counterparts (**Fig. 1f,i**). In contrast, no p-S6 (Ser235/236) difference was found between wild-type and mutant GFP+ cells in arteries. These data show that *Pik3ca^H1047R^*ECs selectively expand in capillaries and veins and that arteries develop normally despite mutant cells.

**Figure 1.**
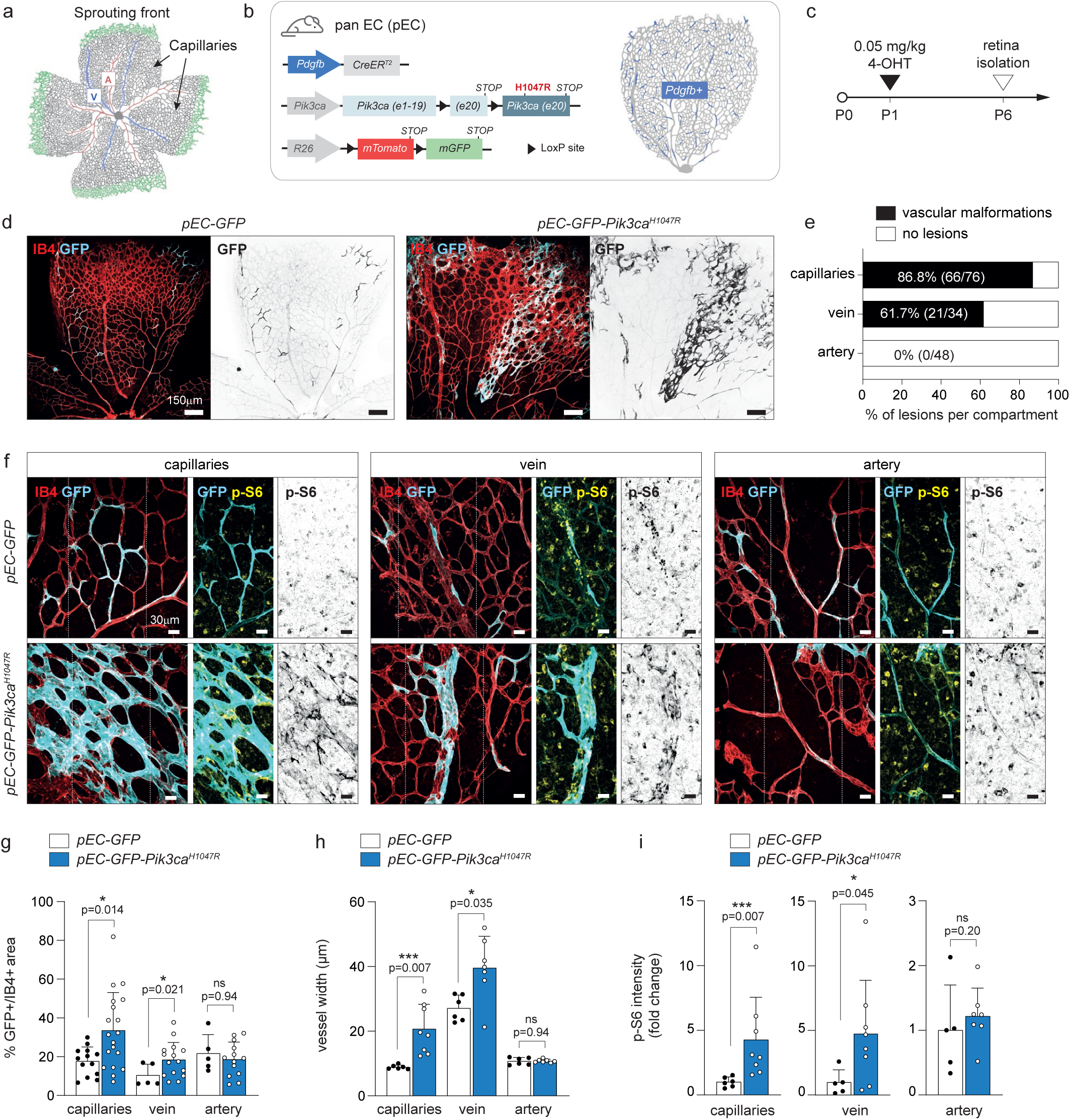
Spatial mapping of *Pik3ca^H1047R^* clone expansions in the retinal vasculature. (a) Schematic illustration of retinal vasculature. Veins (V) are shown in blue, arteries (A) are shown in red, the capillary network is represented in grey and the sprouting front in green. (b) Left: Schematic illustration of the transgenic mouse strategy combining the pan-endothelial (pEC) tamoxifen-inducible Cre (*Pdgfb-CreER^T2^*) allele with either (i) the *R26-mTmG* reporter allele alone (*pEC-GFP*), or (ii) the *R26-mTmG* reporter together with the *Pik3ca^H1047R^* allele (*pEC-GFP-Pik3ca^H1047R^*), both Cre-dependent. Right: Schematic of the growing vasculature in a petal of the retina showing in blue the population targeted by activating *Pdgfb-CreER^T2^* in a stochastic fashion. (c) Experimental timeline showing postnatal administration of 0.05 mg/kg of 4-hydroxytamoxifen (4-OHT) at P1 littermates, followed by retina isolation and analysis at P6. (d) Representative confocal images of pEC-*GFP* and *pEC-GFP-Pik3ca^H1047R^* P6 mouse retinas stained for Isolectin B4 (IB4, red, blood vessels) and GFP (cyan, recombined cells). (e) Quantification of the incidence of vascular malformations in capillaries, veins, and arteries in *pEC-GFP*-*Pik3ca^H1047R^* P6 retinas, expressed as the percentage of capillary beds, veins or arteries exhibiting a pathological overgrowth. (f) High-magnification confocal images of IB4-(red), GFP-(cyan) and phospho (p)-S6 (Ser235/236)-(yellow) stained P6 *pEC-GFP* and *pEC-GFP-Pik3ca^H1047R^* retinas. White dotted rectangles mark regions of interest (ROIs), which are shown in the adjacent right panel. Note the expansion of GFP+ ECs and increased S6 phosphorylation in capillaries and veins but not in arteries in *pEC-GFP-Pik3ca^H1047R^* retinas. (g, h) Quantification of the percentage of GFP+ area (n ≥ 5 retinas per genotype) (g) and vessel width (μm, n ≥ 6 retinas per genotype) (h) in capillaries, veins and arteries from *pEC-GFP* and *pEC-GFP-Pik3ca^H1047R^* P6 retinas. (i) Quantification of p-S6 intensity within GFP+ cells in capillaries, veins, and arteries, expressed as fold change comparing *pEC-GFP* to *pEC-GFP-Pik3ca^H1047R^* vasculature (n ≥ 5 retinas per genotype). Data are presented as mean ± s.d. Statistical analysis was performed using nonparametric Mann-Whitney test. ns, non-statistical. Scale bars: 150 µm (d), 30 µm (f).

### Differentiated arterial cells do not respond to *Pik3ca^H1047R^*

The lack of response to *Pik3ca^H1047R^* by arterial ECs led us to hypothesize that mutant cells could be ejected from arteries over time. To investigate this hypothesis, we used the *Bmx(PAC)-CreER^T2^*mouse line, which targets mature arterial ECs in the retinal vasculature. As described above, wild-type and *Pik3ca^H1047R^* arterial ECs were traced by combining *Pik3ca^H1047R^*; *Bmx(PAC)-CreER^T2^* with *R26-mTmG* (hereafter referred to as *aEC-GFP* and *aEC-GFP-Pik3ca^H1047R^,* respectively; **Fig. 2a**). First, we characterized *Bmx* promoter activity by treating *aEC-GFP* pups with a single saturating dose of 4-OHT at three different postnatal stages (P1, P4, and P7), followed by isolating retinas five days later (P6, P9, and P12, respectively) (**Extended Data Fig. 2a,b**). Inducing *aEC-GFP* mice at P1 did not label any arterial ECs at P6, indicating a lack of mature arteries at the induction stage. Targeting *Bmx+* cells at P4 and P7 resulted in an overt presence of GFP+ cells in arteries (**Extended Data Fig. 2b**). In agreement with *Bmx* being a marker of mature arteries, the number of GFP+ cells was higher when 4-OHT was injected at P7 (**Extended Data Fig. 2b**). Based on these results, we analyzed *aEC-GFP-Pik3ca^H1047R^* retinas by injecting 4-OHT at P4 or P7 (**Fig. 2b-d**). In these retinas, the proportion of GFP+ cells and the width of arteries were similar to those in *aEC-GFP* littermate retinas (**Fig. 2c,d,g,h**). In line with the data shown above (**Fig. 1f,i**), *Pik3ca^H1047R^*-GFP+ arteries did not exhibit an increase in PI3K signaling (**Fig. 2e-h**). Further supporting the lack of pathological phenotypes, arteries of *aEC-GFP* and *aEC-GFP-Pik3ca^H1047R^* retinas showed similar coverage of smooth muscle cells (**Extended Data Fig. 2c**). Next, we studied the long-term effects of *Pik3ca^H1047R^* mutant cells in the arteries by analyzing retinas at P21. We found no differences in the GFP+ area or distribution of wild-type and *Pik3ca^H1047R^*-GFP+ ECs in the main arteries and arteriolar branches (**Fig. 2I, j**). These results confirm that mutant cells are well-tolerated in arteries for long periods without triggering any pathological response.

**Figure 2.**
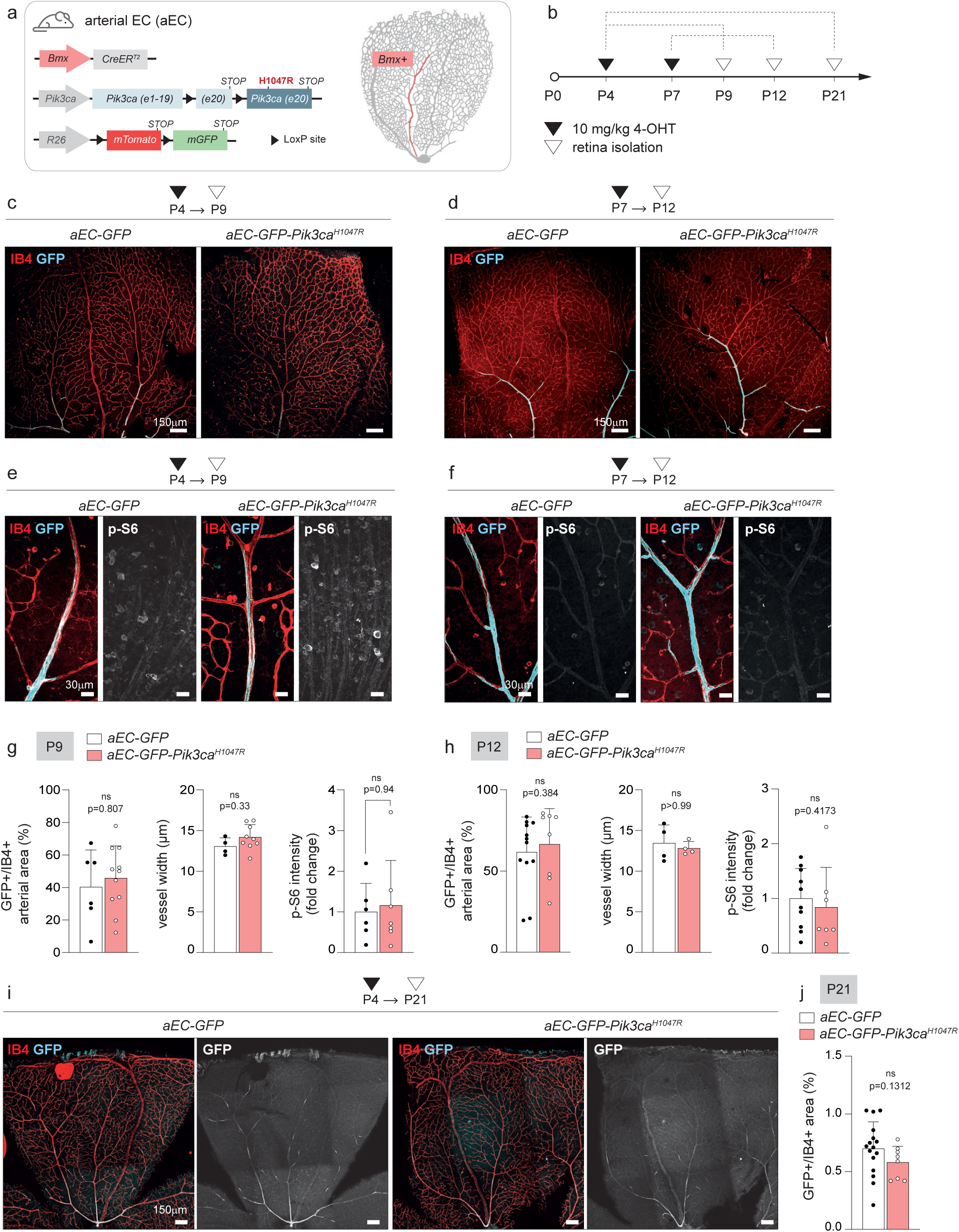
**Differentiated arterial cells do not respond to *Pik3ca^H1047R^*** ((a) Left: Schematic illustration of the transgenic mouse strategy combining the arterial EC-specific (aEC) tamoxifen-inducible Cre (*Bmx(PAC)-CreER^T2^*) allele with either (i) the *R26-mTmG* reporter allele alone (*aEC-GFP*), or (ii) the *R26-mTmG* reporter together with the *Pik3ca^H1047R^* allele (*aEC-GFP*-*Pik3ca^H1047R^*), both Cre-dependent. Right: Schematic of the growing vasculature in a petal of the retina showing in red the population targeted by activating *Bmx(PAC)-CreER^T2^*. (b) Experimental timeline showing postnatal administration of 10mg/kg of 4-OHT at P4 or P7, followed by retina isolation and analysis at P9, P12 or P21. (c,d) Representative confocal images of *aEC-GFP* and *aEC-GFP-Pik3ca^H1047R^* P9 (c) and P12 (d) retinas immunostained for IB4 (red, blood vessels) and GFP (cyan, recombined cells) showing the distribution of GFP+ cells along the arteries. (e,f) High-magnification images of arteries from *aEC-GFP* and *aEC-GFP-Pik3ca^H1047R^* P9 (e) and P12 (f) retinas showing immunodetection of GFP (cyan, recombined cells), phospho (p)-S6 (Ser235/236) (white) and IB4 (red). (g,h) Quantification of the percentage of GFP+ arterial area (left), artery width (middle) and p-S6 (Ser235/236) intensity within GFP+ area (right) comparing *aEC-GFP* and *aEC-GFP*-*Pik3ca^H1047R^* P9 (g) and P12 (h) retinas (n ≥ 4 retinas per genotype). (i) Representative confocal images of *aEC-GFP* and *aEC-GFP*-*Pik3ca^H1047R^* P21 retinas, induced with 10 mg/kg 4-OHT at P4. Retinas were stained for IB4 (red, blood vessels) and GFP (cyan, recombined cells). (j) Quantification of GFP+ area, presented as a percentage of GFP+/IB4+ area comparing *aEC-GFP* and *aEC-GFP*-*Pik3ca^H1047R^* P21 retinas within the total vasculature (n ≥ 8 retinas per genotype). Data are presented as mean ± s.d. Statistical analysis was performed by nonparametric Mann-Whitney test. ns, non-statistical. Scale bars: 150 µm (c,d,i) and 30 µm (e,f).

### *Pik3ca^H1047R^* expression reduces arterial mobilization

To investigate whether the lack of pathological response in mature arteries was a matter of differentiation stage, we next targeted pre-arterial cells. Tip ECs express arterial markers and contribute to expanding arteries (Xu et al. 2014; Pitulescu et al. 2017); thus, they are considered pre-arterial cells. To trace tip ECs and their progeny, we used the *Esm1(BAC)-CreER^T2^*mouse line crossed with the *R26-mTmG* (*tEC-GFP*) alone or in combination with *Pik3ca^H1047R^* (*tEC-GFP-Pik3ca^H1047R^*, **Fig. 3a**). 4-OHT was administrated at P1, followed by lineage tracing of the *Esm1*-GFP+ progeny at P2, P6 and P8 (**Fig. 3b,c**). Twenty-four hours after 4-OHT administration, GFP+ cells were mainly detected at the sprouting front of *tEC-GFP* and *tEC-GFP-Pik3ca^H1047R^* retinas (**Fig. 3c**). At P6, arteries of *tEC-GFP* retinas were extensively composed of wild-type-GFP+ cells, and the GFP+ area in arteries increased by P8 (**Fig. 3c**). Consistent with a proportion of *Esm1+* cells contributing to the capillary bed (Pitulescu et al. 2017), we also detected wild-type-GFP+ cells distributed along capillaries. Surprisingly, we found that the area of GFP+ ECs that covered arteries at P6 and P8 was reduced by half in *tEC-GFP-Pik3ca^H1047R^* mice (**Fig. 3c,e,f**). Remarkably, none of the few *Pik3ca^H1047R^*-GFP+ cells that reached an artery triggered arterial overgrowth (**Fig. 3d**). Most *Esm1*-traced *Pik3ca^H1047R^*-GFP+ cells remained in the capillary bed, substantially expanding (**Fig. 3c,e,f**) and forming multiple vascular malformations (**Fig. 3d**). The expansion of the mutant clones in capillaries was associated with a significant increase in PI3K pathway signaling (**Fig. 3g,h**) and enhanced cell proliferation (**Extended Data Fig. 3a-c**). Intriguingly, many veins in *tEC-GFP-Pik3ca^H1047R^* retinas also contained GFP+ cells, a phenotype rarely seen in control retinas. We also observed that the proportion of GFP+ area in the veins of *tEC-GFP-Pik3ca^H1047R^* mice grew over time (P8 *vs.* P6) (**Fig.3f**). The loss of *Pik3ca^H1047R^*-GFP+ cells in arteries and their gain in veins suggest that *Pik3ca^H1047R^*induces a fate switch. We also analyzed P21 retinas and found that *Pik3ca^H1047R^*-GFP+ clones remained in veins and capillaries (**Extended Data Fig. 3d-f**). Consistent with the expansion of mutant clones in the presence of mitogenic signals(Kobialka et al. 2022), there was no further expansion of *Pik3ca^H1047R^*-GFP+ cells at later time points (P21), in contrast to their active expansion at P6 and P8 (**Extended Data Fig. 3g**). These data indicate that *Pik3ca^H1047R^* expression in arterial precursors prevents their mobilization to arteries, thereby remaining in capillaries and veins where they clonally expand and form vascular malformations.

**Figure 3.**
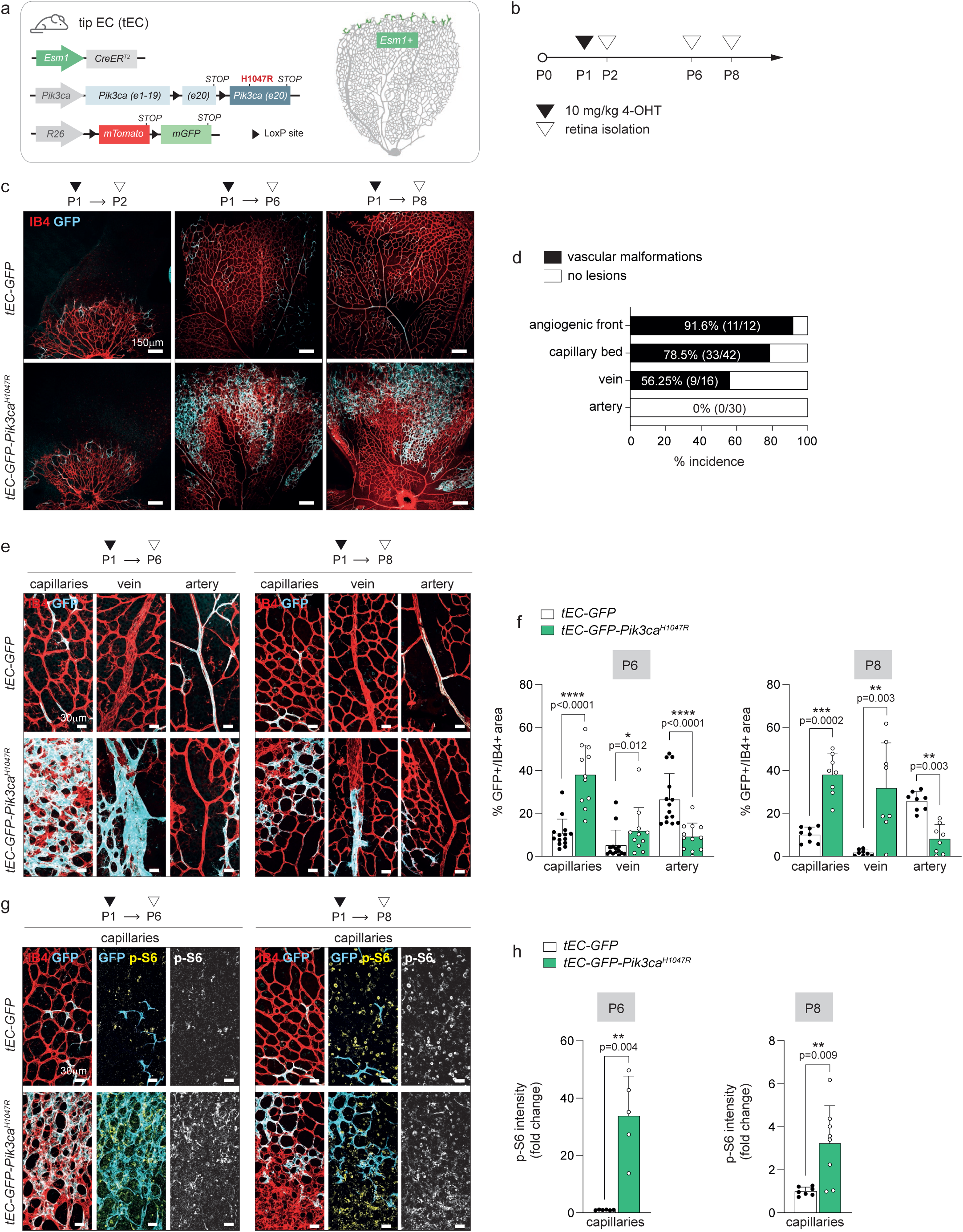
***Pik3ca^H1047R^* expression interferes with arterial mobilization** (a) Left: Schematic illustration of the transgenic mouse strategy combining the tip EC-specific (*tEC*) tamoxifen-inducible Cre (*Esm1(BAC)-CreER^T2^*) allele with either (i) the *R26-mTmG* reporter allele alone (*tEC-GFP*), or (ii) the *R26-mTmG* reporter together with the *Pik3ca^H1047R^* allele (*tEC-GFP*-*Pik3ca^H1047R^*), both Cre-dependent. Right: Schematic of the growing vasculature in a petal of the retina showing in green the population targeted by activating *Esm1(BAC)-CreER^T2^*. (b) Experimental timeline showing postnatal administration of 10 mg/kg of 4-OHT at P1 littermates, followed by retina isolation and analysis at P2, P6, and P8. (c) Representative confocal images of *tEC-GFP* and *tEC-GFP-Pik3ca^H1047R^* of P2, P6 and P8 retinas stained with IB4 (red, blood vessels) and anti-GFP (cyan, recombined cells). (d) Quantification of the incidence of vascular malformations in the sprouting front, capillaries, veins, and arteries of P6 *tEC-GFP*-*Pik3ca^H1047R^* retinal vasculature, expressed as the percentage of vessels or capillary beds exhibiting a pathological phenotype. (e) High-magnification confocal images of *tEC-GFP* and *tEC-GFP-Pik3ca^H1047R^* P6 (left panel) and P8 (right panel) retinas stained with IB4 (red, blood vessels) and anti-GFP (cyan, recombined ECs) showing the expansion of GFP+ ECs (P1 *Esm1*-derived progeny) in capillaries, veins, and arteries. (f) Quantification of GFP+ EC expansion (P1 *Esm1*-derived progeny) in *tEC-GFP* and *tEC-GFP-Pik3ca^H1047R^* P6 (left) and P8 (right) retinas, comparing capillaries, veins, and arteries (n ≥ 8 retinas per genotype). (g) High-magnification confocal images of capillary regions from *tEC-GFP* and *tEC-GFP-Pik3ca^H1047R^* P6 (left panel) and P8 (right panel) retinas, stained for IB4 (red, blood vessels), GFP (cyan, recombined cells), and phospho (p)-S6 (Ser235/236) (yellow). (h) Quantification of p-S6 (Ser235/236) intensity within the GFP+ capillary area comparing *tEC-GFP* and *tEC-GFP*-*Pik3ca^H1047R^* P6 (left) and P8 (right) retinas (n ≥ 5 retinas per genotype). Data are presented as mean ± s.d. Statistical analysis was performed using nonparametric Mann-Whitney test. Scale bars: 150 µm (c) and 30 µm (e, g).

### Multispectral mosaic tracing of wild-type and *Pik3ca^H1047R^* endothelial clones

To unequivocally trace both wild-type and mutant cells in the same tissue microenvironment, we generated a new genetic mouse model, hereafter abbreviated as *_iFluMosaic-PIK3CA_H1047R* _(*Rosa26*_*CAG-LSL-H2B-Cherry/H2B-GFP-2A-HAHuPIK3CAH1047R*). This new allele contains two pairs of distinct and mutually exclusive Lox sites, which Cre can stochastically recombine to induce the expression of either *His-H2B-Cherry* or equimolar co-expression of *H2B-EGFP-V5* along with *HA*-*PIK3CA^H1047R^*, due to the viral 2A peptide separating them (**Extended Data Fig. 4a**). First, we assessed the functionality of the construct by breeding *iFluMosaic-PIK3CA**^H1047R^***with *Cdh5(PAC)-CreER^T2^* mice (**Extended Data Fig. 5a**). Lung ECs from these mice were isolated, cultured, and treated with 4-OHT to induce stochastic recombination of the mutually exclusive LoxP sites (**Extended Data Fig. 5b**). Upon early recombination, we observed that the proportion of Cherry+ ECs (wild type, ≈60%) was higher than EGFP-*PIK3CA*^H1047R^ ECs (≈40%) (**Extended Data Fig. 5c,e**). These data are consistent with the notion that the LoxN sites in front of the Cherry reporter are closer together than the Lox2272 sites and follow the ratiometric recombination pattern shown in similar constructs (Pontes-Quero et al. 2017). As expected, EGFP-*PIK3CA^H1047R^* ECs showed higher proliferative rates (**Extended Data Fig. 5c,f**) and higher PI3K signaling than Cherry-Wild-type ECs (**Extended Data Fig. 5d,g**). This higher proliferation led to a higher fraction of EGFP-*PIK3CA^H1047R^* ECs over time, offsetting the initial Cherry bias (**Extended Data Fig. 5c-e**).

Next, we fate-mapped EGFP-*PIK3CA^H1047R^* and Cherry-Wild-type pre-arterial tip ECs *in vivo* by crossing *iFluMosaic-PIK3CA^H1047R^* with *Esm1(BAC)-CreER^T2^* mice (*tEC-iFluMosaic-PIK3CA^H1047R^*). Pups were treated with a high dose of 4-OHT at P1 and P2, followed by assessing the behavior of EGFP-*PIK3CA^H1047R^*and Cherry-Wild-type targeted cells 6 days after (**Fig. 4a,b**). Cherry+ cells were distributed between capillaries and arteries (**Fig. 4c,d**). In contrast, *Esm1*-targeted EGFP-*PIK3CA^H1047R^* cells and their progeny were found in veins and capillaries of the sprouting front, the central plexus, and the immediate vicinity of arteries (**Fig. 4c,d**). In these locations, EGFP-*PIK3CA^H1047R^* cells clonally expanded and formed vascular malformations (**Fig. 4d**), which overall resulted in a more significant number of EGFP+ cells (i.e., 7680) compared to Cherry+ cells (i.e., 511) (**Fig. 4e**). Vascular lesions were of different sizes, including big (> 30 cells), medium (between 10-20 cells), and small (<10 cells) (**Fig. 4f**). EGFP-*PIK3CA^H1047R^* cells showed increased Ki67 positivity (**Fig. 4g** and **Extended Data Fig. 5h**) and p-S6 immunostaining compared to Cherry-Wild-type cells (**Fig. 4h** and **Extended Data Fig. 5i**). We noticed that few EGFP-*PIK3CA^H1047R^* cells (0,47% of total EGFP+ cells) settled in the artery, albeit at an almost fifty-fold lower proportion than Cherry+ cells (23% of total Cherry+ cells) (**Fig. 4d, e**). Overall, among arteries that contain recombined cells only 16% presented EGFP-*PIK3CA^H1047R^*mutant ECs (**Fig. 4i**). EGFP-*PIK3CA^H1047R^* cells that arrived at the artery formed small lesions (composed of an average of 7 cells/lesion) (**Fig. 4d,j**). Given that malformations in the artery do not occur when expression of *Pik3ca^H1047R^* is regulated through the endogenous locus (**Fig. 3**), these data suggest that pre-arterial cells downregulate the levels of PI3Kα before differentiating into mature arterial cells. Next, we wondered whether similar phenotypes were observed upon forced overexpression of *PIK3CA^H1047R^* in mature arterial cells. We combined *iFluMosaicPIK3CA^H1047R^* with *Bmx(PAC)-CreER^T2^*and injected 4-OHT between P7 and P9 (**Fig. 4k,l**). P12 arteries contained the expected ratio of Cherry+ (65%) and EGFP-*PIK3CA^H1047R^* (35%) cells (**Fig. 4m,n**). Yet, mutant EGFP+ cells in mature arterial ECs did not trigger any pathological phenotype (**Fig. 4o**), indicating that forced overexpression of *PIK3CA^H1047R^* in mature arterial cells is insufficient to overcome their resistance to expand upon expression of these mutations.

**Figure 4.**
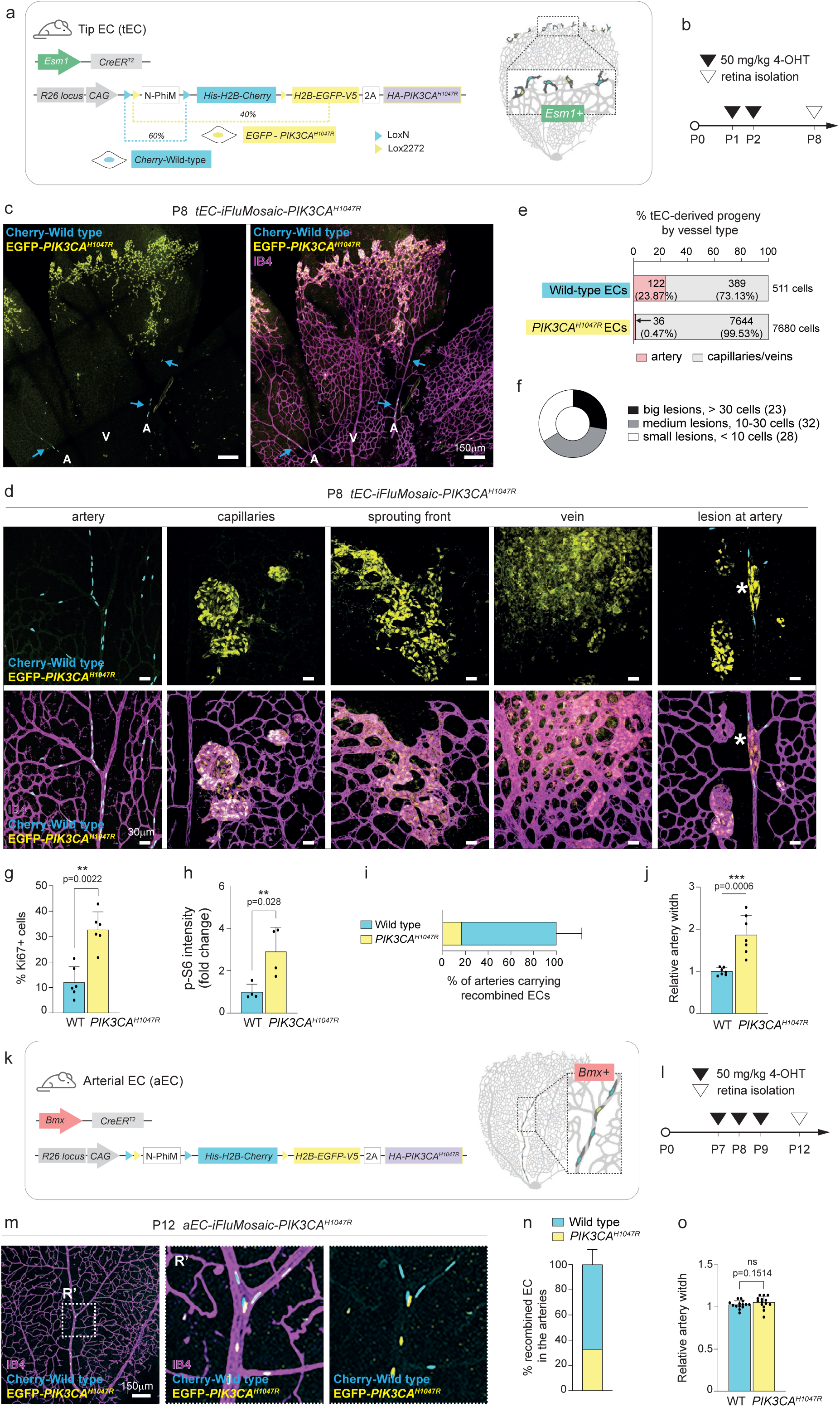
Multispectral mosaic tracing of wild-type and *Pik3ca^H1047R^* endothelial clones. (a) Left: Schematic illustration of the transgenic mouse strategy combining the tip EC-specific (*tEC*) tamoxifen-inducible Cre (*Esm1(BAC)*-CreER^T2^) allele with the *iFluMosaic-PIK3CA^H1047R^* allele. The *iFluMosaic-PIK3CA^H1047R^* knock-in model contains two mutually exclusive *Lox* sites that, upon recombination by Cre activity, either result in the expression of (i) the nuclear His-H2B-Cherry (recombined cell remains wild type for *Pik3ca*) or (ii) H2B-EGFP-V5 and HA-*PIK3CA^H1047R^* separated by a 2A peptide (recombined cell overexpresses human *PIK3CA^H1047R^* cDNA tagged with HA at the C-terminal along with the nuclear EGFP). This system allows for the simultaneous tracing of both wild-type and mutant cells within the same sample using coupled fluorescent proteins. For detailed information, refer to Figure S4 and M&M section. Right: Schematic of the growing vasculature in a petal of the retina showing in dark grey the population targeted by activating *Esm1(BAC)-CreER^T2^*. Recombined cells are distinguished by fluorescent nuclear markers: Cherry-Wild-type EC for *Pik3ca* are represented in cyan, while ECs expressing EGFP and *PIK3CA^H1047R^* are represented in yellow. (b) Experimental timeline showing postnatal administration of 50 mg/kg of 4-OHT at P1 and P2 mouse littermates, followed by retina isolation and analysis at P8. (c) Representative confocal images of *tEC-iFluMosaic-PIK3CA^H1047R^* P8 mouse retinas immunostained for IB4 (magenta, blood vessels), Cherry (cyan, wild-type ECs for *Pik3ca*) and EGFP (yellow, *PIK3CA^H1047R^* expressing ECs). “A” refers to the artery, “V” refers to the vein. Cyan arrows point to wild-type ECs for *Pik3ca*. (d) High-magnification confocal images of *tEC-iFluMosaic-PIK3CA^H1047R^* P8 retinas. From left to right: healthy artery populated with Cherry-Wild-type ECs, malformed capillaries, sprouting front, and vein, showing pathological expansion of EGFP-*PIK3CA^H1047R^* clones, and a malformed artery (indicated with a white asterisk) with EGFP-*PIK3CA^H1047R^* clones, as an example of an occasional lesion. (e) Stacked bar graph comparing the percentage of P1 and P2 recombined *Esm1*-Cherry-Wild-type and EGFP-*PIK3CA^H1047R^*-derived progeny located within arteries *vs* capillaries and veins in P8 *tEC-iFluMosaic-PIK3CA^H1047R^* retinas. Numbers within the bar show the total number of analyzed cells and the resulting percentage for each category (n = 6 retinas). (f) Pie chart categorizing lesions based on their extent, defined by the number of EGFP-*PIK3CA^H1047R^* ECs in *tEC-iFluMosaic-PIK3CA^H1047R^* P8 retinas (n = 14 retinas). (g) Quantification of cell proliferation assessed as a percentage of Ki67+ staining within Cherry-Wild-type and EGFP-*PIK3CA^H1047R^* ECs in *tEC-iFluMosaic-PIK3CA^H1047R^* P8 retinas (n = 6 retinas). (h) Quantification of phospho (p)-S6 (Ser235/236) intensity in Cherry-Wild-type or EGFP-*PIK3CA^H1047R^* ECs of *tEC-iFluMosaic-PIK3CA^H1047R^* P8 retinas (n = 4 retinas). (i) Quantification of the percentage of arteries containing Cherry-Wild-type and EGFP-*PIK3CA^H1047R^* ECs among total number of arteries with any recombined EC. (j) Quantification of artery width in regions containing Cherry-Wild-type and EGFP-*PIK3CA^H1047R^* ECs compared to areas with non-labeled wild-type ECs (n = 7 arteries from 6 retinas). This parameter is used to detect arterial overgrowth. (k) Left: Schematic illustration of the transgenic mouse strategy combining the artery EC-specific (aEC) tamoxifen-inducible Cre (*Bmx(PAC)*-CreER^T2^) allele with *iFluMosaic-PIK3CA^H1047R^* allele. Right: Schematic of the growing vasculature in a petal of the retina showing in dark grey the population targeted by activating *Bmx(PAC)*-CreER^T2^. Recombined cells are distinguished by fluorescent nuclear markers: Cherry-Wild-type EC for *Pik3ca* are represented in cyan, while ECs expressing EGFP and *PIK3CA^H1047R^* are represented in yellow. (l) Experimental timeline showing postnatal administration of 50mg/kg of 4-OHT at P7, P8 and P9 mouse littermates, followed by retina isolation and analysis at P12. (m) A representative confocal image of *aEC-iFluMosaic-PIK3CA^H1047R^*P12 mouse retina immunostained for IB4 (magenta, blood vessels), Cherry (cyan, wild-type ECs for *Pik3ca*) and EGFP (yellow, *PIK3CA^H1047R^* expressing ECs). The white dotted square marks the ROI (R’), which is magnified to show an artery in greater detail in the right panel. (n) Quantification of the percentage of recombined Cherry-Wild-type and EGFP-*PIK3CA^H1047R^* ECs within *aEC*-*iFluMosaic-PIK3CA^H1047R^* P12 arteries (n = 14 retinas). (o) Quantification of artery width in regions containing Cherry-Wild-type and EGFP-*PIK3CA^H1047R^* ECs compared to regions with non-labeled wild-type ECs (n= 14 retinas). Data are presented as mean ± s.d. Statistical analysis was performed using nonparametric Mann-Whitney test (g,h,j, and o). ns, non-statistical. Scale bars: 150 µm (c, m) and 30 µm (d).

Our data demonstrate that arterial differentiation is interrupted upon acquiring *PIK3CA^H1047R^*mutations in pre-arterial cells.

### *Pik3ca^H1047R^* expression triggers a transcriptional switch toward a venous phenotype

To study how *Pik3ca* mutations induce a cell fate switch, we analyzed the transcriptomic profile of ECs by single-cell RNA sequencing (scRNAseq). Given the low number of *Esm1*+ cells in the retina, particularly in wild-type mice, we chose an *in vitro* strategy. We aimed to capture the presence of various EC subsets and identify whether acute expression of *Pik3ca^H1047R^* interfered with cell fate and state transitions. To study these questions, we used ECs derived from *Pdgfb-CreER^T2^; Pik3ca^H1047R^* (*pEC-Pik3ca^H1047R^*) mice. Parental ECs were treated with vehicle (wild type for *Pik3ca*) or 4-OHT (expressing the *Pik3ca^H1047R^*allele) and analyzed at 48 h post-induction. scRNAseq analysis of control and *Pik3ca^H1047R^* ECs yielded 5 clusters (C1-C5) with different signatures of marker enrichment in the two-dimensional (2D) Uniform Manifold Approximation and Projection (UMAP) (**Fig. 5a** and **Extended Data Fig. 6a,b**). Specifically, the C1 cluster exhibited expression of genes characteristic of tip and arterial ECs, such as *Cxcr4, Esm1, Apln,* and *Sox17* (**Fig. 5d** and **Extended Data Fig. 6b**). The C2 cluster contained cells with high expression of known markers of venous identity, including *Nrp2, Ephb4,* and *Nr2f2*. Some cells within the C2 cluster also showed enrichment for canonical markers of lymphatic ECs (*Prox1, Lyve-1, Itga9*) (**Extended Data Fig. 6c**). The C4 and C5 clusters showed enrichment of genes involved in cell proliferation, while the C3 cluster contained cells with elevated expression of inflammation-related genes (*Cd44, Ptgs2, Serpine1, Cxcl12,* **Fig. 5a,d** and **Extended Data Fig. 6b**).

**Figure 5.**
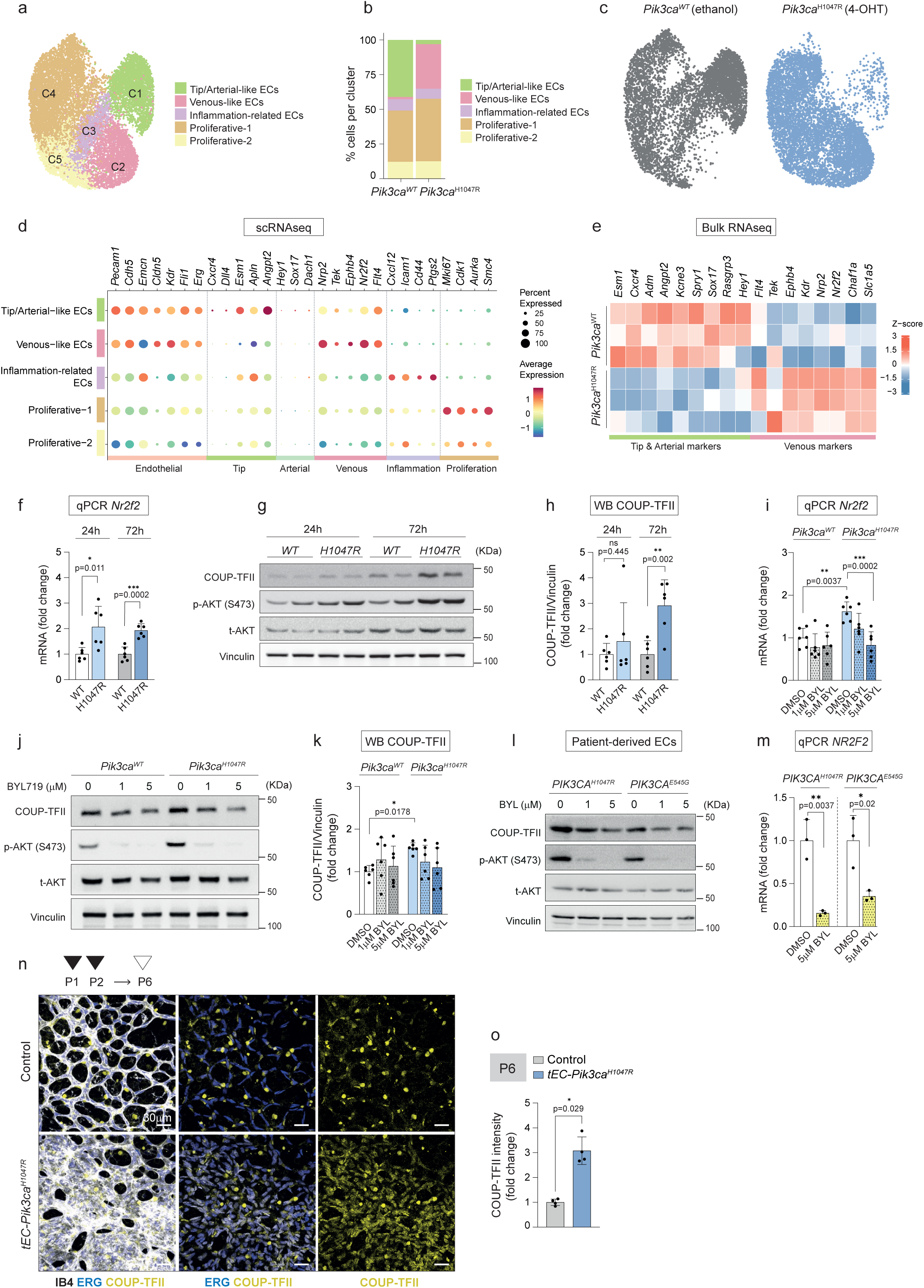
***Pik3ca^H1047R^* expression promotes a transcriptional shift towards a venous phenotype with an increase in *Nr2f2* levels** (a) UMAP representation of Louvain clustering of merged *Pdgfb-CreER^T2^* (pEC)-*Pik3ca^H1047R^* mouse lung endothelial cells (ECs) 48 h after treatment with ethanol (vehicle, wild type for *Pik3ca*) and 4-OHT (expressing *Pik3ca^H1047R^*) derived from scRNA-seq data. (b) Bar plot showing the distribution of *pEC*-Wild-type (ethanol-treated) and *pEC-Pik3ca^H1047R^* (4-OHT-treated) ECs clusters. (c) UMAP representation of *pEC-*Wild-type (ethanol-treated, grey) and *pEC-Pik3ca^H1047R^* (4-OHT-treated, blue) ECs shown independently. (d) Dot plot showing the average expression of key cell type markers across the 5 clusters. (e) Tip, arterial and venous markers selected from differentially expressed genes (DEG, FDR adjusted p-value < 0.05) analysis in bulk RNA-seq data (Kobialka et al. 2022) comparing *pEC-Pik3ca^H1047R^* (4-OHT-treated) with *pEC*-Wild-type (ethanol-treated) ECs 24 h post-induction. (f) RT-qPCR for *Nr2f2* gene expression 24 h and 72 h after ethanol or 4-OHT treatment in *pEC-Pik3ca^H1047R^* ECs (n = 6 biological replicates). (g) Representative immunoblot showing the activation of PI3K/AKT pathway (by assessing the levels of phopsho(p)-AKT (Ser473)) and COUP-TFII protein levels in *pEC-Pik3ca^H1047R^* ECs 24 h and 72 h after ethanol or 4-OHT treatment. (h) Quantification of COUP-TFII protein levels normalized to vinculin levels (n = 6 biological replicates). (i) RT-qPCR for *Nr2f2* gene expression in *pEC-Pik3ca^H1047R^* ECs treated with the PI3Kα specific inhibitor, BYL719. ECs were first treated with ethanol or 4-OHT for 6 h, followed by treatment with vehicle or BYL719 (1 µM and 5 µM) for 66 h. (n = 6 biological replicates). (j) Representative immunoblot showing the impact BYL719 inhibitor on PI3K/AKT signaling and COUP-TFII levels in *pEC-Pik3ca^H1047R^* ECs. ECs were first treated with ethanol or 4-OHT for 6 h, followed by treatment with vehicle or BYL719 (1 µM and 5 µM) for 66 h. (k) Quantification of COUP-TFII protein levels normalized to vinculin levels in *pEC-Pik3ca^H1047R^* ECs treated with BYL719 (vehicle DMSO, 1 µM and 5 µM) (n = 6 biological replicates). (l) Representative immunoblot showing the impact of BYL719, on PI3K/AKT signaling and COUP-TFII levels in *PIK3CA^H1047R^* and *PIK3CA^E545G^* patient-derived venous ECs. These ECs were treated with vehicle DMSO and BYL719 (1 µM and 5 µM) for 72 h. (m) RT-qPCR for *NR2F2* gene expression in *PIK3CA^H1047R^* and *PIK3CA^E545G^* patient-derived venous ECs treated with BYL719 (vehicle DMSO and 5 µM) (n = 3 technical replicates). (N) Representative confocal images of control and *tEC-Pik3ca^H1047R^* P6 mouse retinas after 25 mg/kg 4-OHT induction at P1 and P2. Immunostaining for IB4 (white, blood vessels), ERG (blue, EC nuclei) and COUP-TFII (yellow) showing upregulation of COUP-TFII in the capillary area of *tEC-Pik3ca^H1047R^* P6 retinas. (o) Quantification of COUP-TFII intensity within ERG+ areas (EC nuclei) in capillary vessels from control and *tEC-Pik3ca^H1047R^* P6 retinas (n = 4 retinas per genotype). Data are presented as mean ± s.d. Statistical analysis was performed using an unpaired t-test (f, h, m), a 2-way ANOVA (i, k), and a nonparametric Mann-Whitney test (o). ns, non-statistical. Scale bars: 30 µm (n).

Consistent with published data (Kobialka et al. 2022), the C4 cluster, composed of proliferative cells, increased by about 10% following *Pik3ca^H1047R^* expression (**Fig. 5b**). We also noticed that there was a marked loss of cells in the C1 cluster enriched in the tip and arterial markers and an overt increase of the C2 venous-like cluster in the 4-OHT treated population (**Fig. 5b,c**). The Principal component analysis of the first two components revaled a high degree of similarity between the C1 and C2 culsters. Hence, we proceed to analyze the differentialy expressed genes (DEG) in these group of cells between *Pik3ca^WT^* and *Pik3ca^H1047R^*-expressing cells (**Extended Data Fig. 6d, e**). This comparison identified that 4141 genes were differentially expressed (FDR adjusted p-value < 0.05), with tip ECs markers strongly decreased (i.e., *Esm1*, *Angpt2, Cxcr4*, *Mecom*, *Adm,* and *Apln*) and venous markers (i.e., *Nrp2*, *Nr2f2,* and *Ephb4)* increased upon expression of *Pik3ca^H1047R^* (**Extended Data Fig. 6f,g**). To validate these molecular changes in an independent data set, we used published bulk RNAseq data (Kobialka et al. 2022). We specifically focused on tip, arterial, and venous genes and found a significantly decreased expression of tip and arterial markers in *Pik3ca^H1047R^* ECs and a prominent increase in expression of venous markers (**Fig. 5e**).

Collectively, these data indicate that in addition to adopting proliferative behavior (Kobialka et al. 2022), expression of *Pik3ca^H1047R^*in ECs induces a molecular transition from tip/arterial cell-like to venous identity phenotype, which aligns with the phenotypes observed in the retinal vasculature.

### *Pik3ca^H0147R^* mutation increases *NR2F2* and COUP-TFII expression

Our scRNAseq and bulk RNAseq data showed that *Pik3ca^H1047R^* expression significantly increased *Nr2f2* mRNA levels (**Extended Data Fig. 6f,g** and **Fig. 5e**). *NR2F2* encodes for COUP-TFII, a key transcription factor (TF) that induces EC proliferation and suppresses arterial identity (Chen et al. 2012; Aranguren et al. 2013; You et al. 2005; Su et al. 2018). Building on these results, we hypothesized that COUP-TFII could act downstream of *Pik3ca^H1047R^* to rewire arterial identity and promote clone expansion. First, we confirmed by quantitative (q)PCR that *Nr2f2* levels rose rapidly 24 h after 4-OHT treatment of mutant ECs *in vitro* and remained elevated over time (**Fig. 5f**). Higher *Nr2f2* mRNA levels in *Pik3ca^H1047R^* ECs led to a significant increase in COUP-TFII protein levels at 72 h post-recombination (**Fig. 5g,h**). PI3K signaling was also overactivated at 24 h and 72 h after 4-OHT treatment, as shown by increased phospho (p)-AKT (Ser473) levels (**Fig. 5g** and **Extended Data Fig. 6h**). Next, we treated these cells with Alpelisib (BYL-719). This PI3Kα-selective inhibitor led to a significant reduction in the levels of p-AKT (Ser473) (**Fig. 5j** and **Extended Data Fig. 6i**) and *Nr2f2* mRNA and COUP-TFII protein in a dose-dependent manner (**Fig. 5i-k**). PI3Kα overactivation also regulated *Nr2f2* and COUP-TFII levels in patient-derived ECs with an activating *PIK3CA* mutation (**Fig. 5l,m**). Finally, COUP-TFII immunostaining of retinas from *tEC-Pik3ca^H1047R^* and control P6 littermates revealed a significant increase in COUP-TFII expression in the progeny of *Esm1*-targeted *Pik3ca^H1047R^* cells *in vivo* (**Fig. 5n,o**). These results establish COUP-TFII as a downstream target of PI3K signaling and that this regulation primarily occurs at the mRNA level.

### *Nr2f2* deletion allows arterialization of *Pik3ca^H1047R^* pre-arterial cells

To elucidate whether COUP-TFII mediates the *Pik3ca^H1047R^*-related fate switch and pathogenic proliferative response *in vivo*, we examined the behavior of *PIK3CA^H1047R^* pre-arterial tip cells in the absence of COUP-TFII expression. For this, we used both genetic strategies of *Pik3ca^H1047R^* expression: endogenous expression, which provides a pathophysiological setting (**Fig. 6a**), and *iFluMosaic-PIK3CA^H1047R^* to faithfully trace wild-type and mutant clones, both in combination with the *Nr2f2^flox/flox^* allele (**Fig. 6f**). To maximize the recombination of all alleles in their endogenous locus, two consecutive doses of 4-OHT were injected (**Fig. 6a,b**). Consequently, more recombined cells were observed in the *tEC-GFP* control retinas (**Fig. 6c**) than those treated with one 4-OHT dose (**Fig. 3c**). First, we confirmed that COUP-TFII expression was reduced in *tEC-Pik3ca^H1047R^-Nr2f2^ΔEC/ΔEC^* retinas (**Extended Data Fig. 7a,b**). Deletion of *Nr2f2* in a wild-type background did not interfere with the mobilization of *Esm1-*targeted *Nr2f2 ^ΔEC/ΔEC^-GFP+* ECs, nor with the overall vascular density (**Fig. 6c,d** and **Extended Data Fig. 7c,d**). *Pik3ca^H1047R^*-*Nr2f2^ΔEC/ΔEC^*-GFP+ mutant cells kept expanding regardless of *Nr2f2* deletion (**Fig. 6c,e**). In contrast, we noticed that *Esm1-*targeted *Pik3ca^H1047R^-Nr2f2^ΔEC/ΔEC^*-GFP+ cells regained the ability to form arteries compared to *Pik3ca^H1047R^-*GFP+ cells (**Fig. 6c,d**). The *tEC*-*iFluMosaic-PIK3CA^H1047R^* model (**Fig. 6f,g**) also showed that EGFP-*PIK3CA^H1047R^-Nr2f2^ΔEC/ΔEC^* mutant cells overexpanded in the capillary bed (**Extended Data Fig. 7e,f**). Yet, while Cherry and Cherry-*Nr2f2^ΔEC/ΔEC^* ECs colonized arteries to a similar extend, EGFP-*PIK3CA^H1047R^-Nr2f2^ΔEC/ΔEC^* cells reached arteries significantly more frequently than EGFP-*PIK3CA^H1047R^* cells (35% *vs* 10% of arteries respectively) (**Fig. 6h,i**). Notably, the majority of EGFP-*PIK3CA^H1047R^-Nr2f2^ΔEC/ΔEC^* cells did not form a lesion when settled in the artery in contrast to EGFP-*PIK3CA^H1047R^*cells, which consistently formed small lesions (**Fig. 6h,j**). These data suggest that tip EGFP-*PIK3CA^H1047R^-Nr2f2^ΔEC/ΔEC^* cells recover the capacity to colonize arteries and differentiate into mature arterial cells despite expressing *PIK3CA^H1047R^*. Collectively, these findings indicate that COUP-TFII orchestrates *PIK3CA*-driven fate switch toward vein identity.

**Figure 6.**
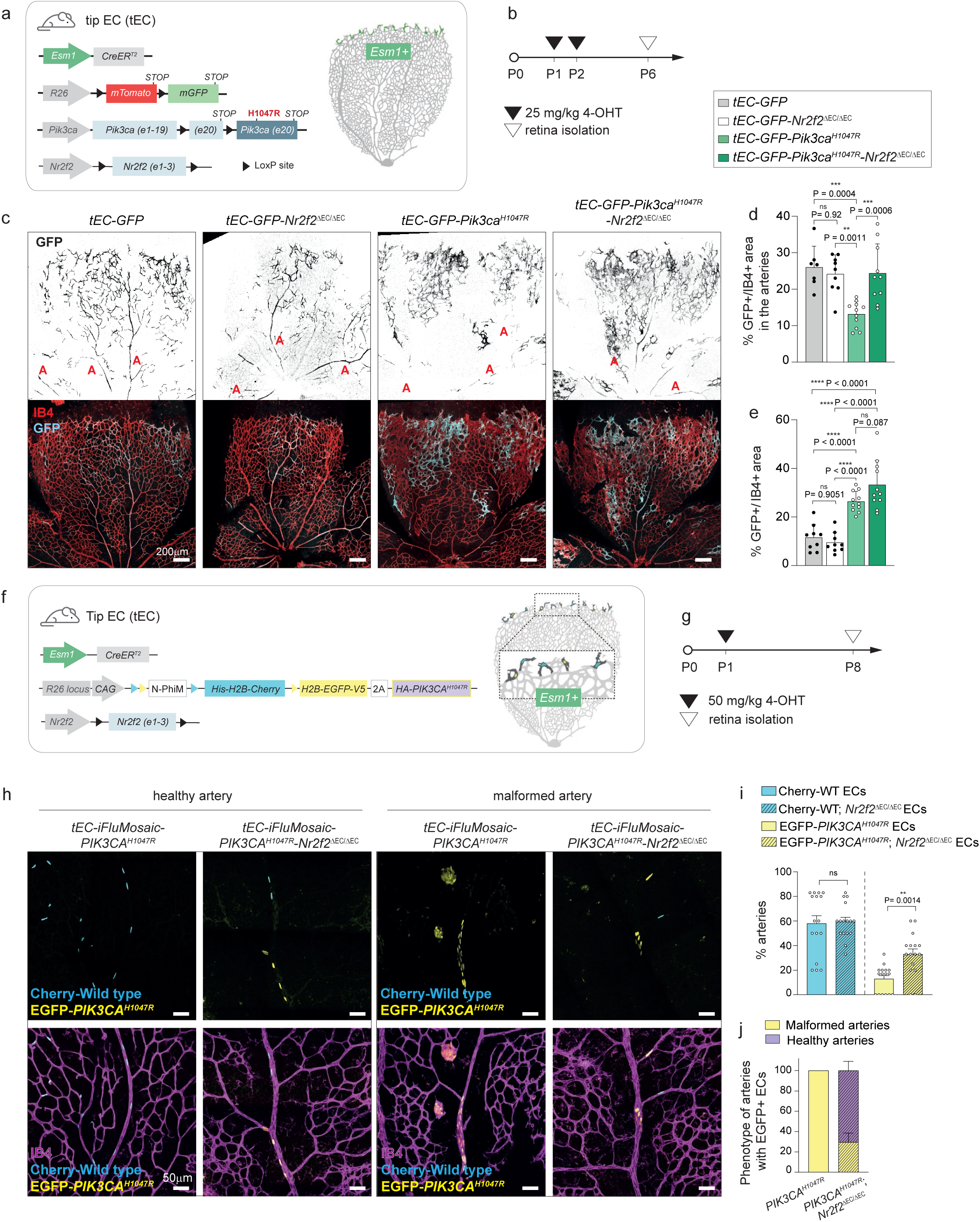
***Nr2f2* deletion allows arterialization of *Pik3ca*^H1047R^ pre-committed arterial cells** (a) Left: Schematic illustration of the transgenic mouse strategy combining the tip EC-specific (tEC) tamoxifen-inducible Cre (*Esm1(BAC)-CreER^T2^*) with the *R26-mTmG* reporter allele (i) alone (*tEC-GFP*), (ii) combined with the *Pik3ca^H1047R^* allele (*tEC-GFP*-*Pik3ca^H1047R^*), (iii) combined with the *Nr2f2*^flox/flox^ allele (*tEC-GFP-Nr2f2*^flox/flox^) or (iv) with both of them (*tEC-GFP-Pik3ca^H1047R^*-*Nr2f2*^flox/flox^). Right: Schematic of the growing vasculature in a petal of the retina showing in green the population targeted by activating *Esm1(BAC)-CreER^T2^*. (b) Experimental timeline showing postnatal administration of 25 mg/kg of 4-OHT at P1 and P2, followed by retina isolation and analysis at P6. (c) Representative confocal images of *tEC-GFP*, *tEC-GFP-Nr2f2^ΔEC/ΔEC^*, *tEC-GFP-Pik3ca^H1047R^*, and *tEC-GFP-Pik3ca^H1047R^*-*Nr2f2^ΔEC/ΔEC^* of P6 mouse retinas, stained for IB4 (red, blood vessels) and GFP (cyan, P1/P2 *Esm1*-derived progeny). “A” denotes for artery. (d, e) Quantification of the GFP+ area specifically within arteries (d) and across all retina vasculature (e), presented as a percentage of the GFP+/IB4+ area, used to assess the expansion of the *Esm1*-derived progeny at P6 (n ≥ 7 retinas per genotype). (f) Left: Schematic illustration of the transgenic mouse strategy combining the tip EC-specific (tEC) tamoxifen-inducible Cre (*Esm1(BAC)-CreER^T2^*) allele with *iFluMosaic-PIK3CA^H1047R^*, either alone or in combination with the *Nr2f2*^flox/flox^ allele. Right: Schematic of the vasculature in a retinal petal showing in dark grey the population targeted by activating *Esm1(BAC)-CreER^T2^*. Recombined cells are distinguished by fluorescent nuclear markers: Cherry-Wild-type EC for Pik3ca are represented in cyan, while ECs expressing EGFP and *PIK3CA^H1047R^* are represented in yellow. (g) Experimental timeline showing postnatal administration of 50 mg/kg of 4-OHT at P1, followed by retina isolation and analysis at P8. (h) Representative confocal images of *tEC-iFluMosaic-PIK3CA^H1047R^* and *tEC-iFluMosaic-PIK3CA^H1047R^-Nr2f2^ΔEC/ΔEC^* P8 retinas stained for IB4 (magenta, blood vessels), Cherry (cyan, wild-type ECs for *Pik3ca*) and EGFP (yellow, *PIK3CA^H1047R^*expressing ECs) in arteries with no lesions (left) and in malformed arteries (right). (i) Quantification of the number of arteries with Cherry+ or EGFP+ ECs, expressed as the percentage of positive arteries out of all arteries in *tEC-iFluMosaic-PIK3CA^H1047R^* and *tEC-iFluMosaic-PIK3CA^H1047R^-Nr2f2^ΔEC/ΔEC^* P8 retinas (n = 16 retinas per genotype). (j) Quantification of EGFP+ arteries in *tEC-iFluMosaic-PIK3CA^H1047R^* and *tEC-iFluMosaic-PIK3CA^H1047R^-Nr2f2^ΔEC/ΔEC^* P8 retinas, categorizing them as healthy or malformed (n ≥ 7 retinas per genotype). Data are presented as mean ± s.d (d, e) or ± s.e.m (i, j). Statistical analysis was performed using one-way ANOVA, followed by Tukey test for multiple comparisons (d, e), and a nonparametric Mann-Whitney test (i). Scale bars: 200 µm (c) and 50 µm (h).

## Discussion

Mutations traditionally associated with cancer have been identified in a variety of diseases, including congenital syndromes, clonal hematopoiesis, and also in healthy tissues. This has sparked the interest of the scientific community in understanding the role of these mutations beyond cancer. In this regard, the impact of these mutations in the epithelium has been extensively studied, providing light on the mechanisms of mutant clone survival (Ciwinska et al. 2024; Herms et al. 2024; Colom et al. 2021). Yet, a question remains: why pathogenic phenotypes are restricted to some tissues and cell types? In this study, we take advantage of the vasculature, a mesoderm-derived tissue commonly affected by cancer mutations, which exhibits restricted pathogenic patterns to these mutations. We specifically focus on *PIK3CA^H1047R^*, a cancer mutation frequently found in capillary, vein, and lymphatic malformations but not in arteries. Our findings identify a wide range of context-dependent effects of *PIK3CA^H1047R^*mutations in ECs, including (i) generation of large clone expansions (i.e., in venous and capillary ECs) causing vascular malformations, (ii) silent presence in refractory cell types (i.e., arterial ECs) without pathological manifestations; and (iii) rewiring of cell fate (i.e., in arterial precursors) favoring pathogenic expansion (**Fig. 7**).

**Figure 7.**
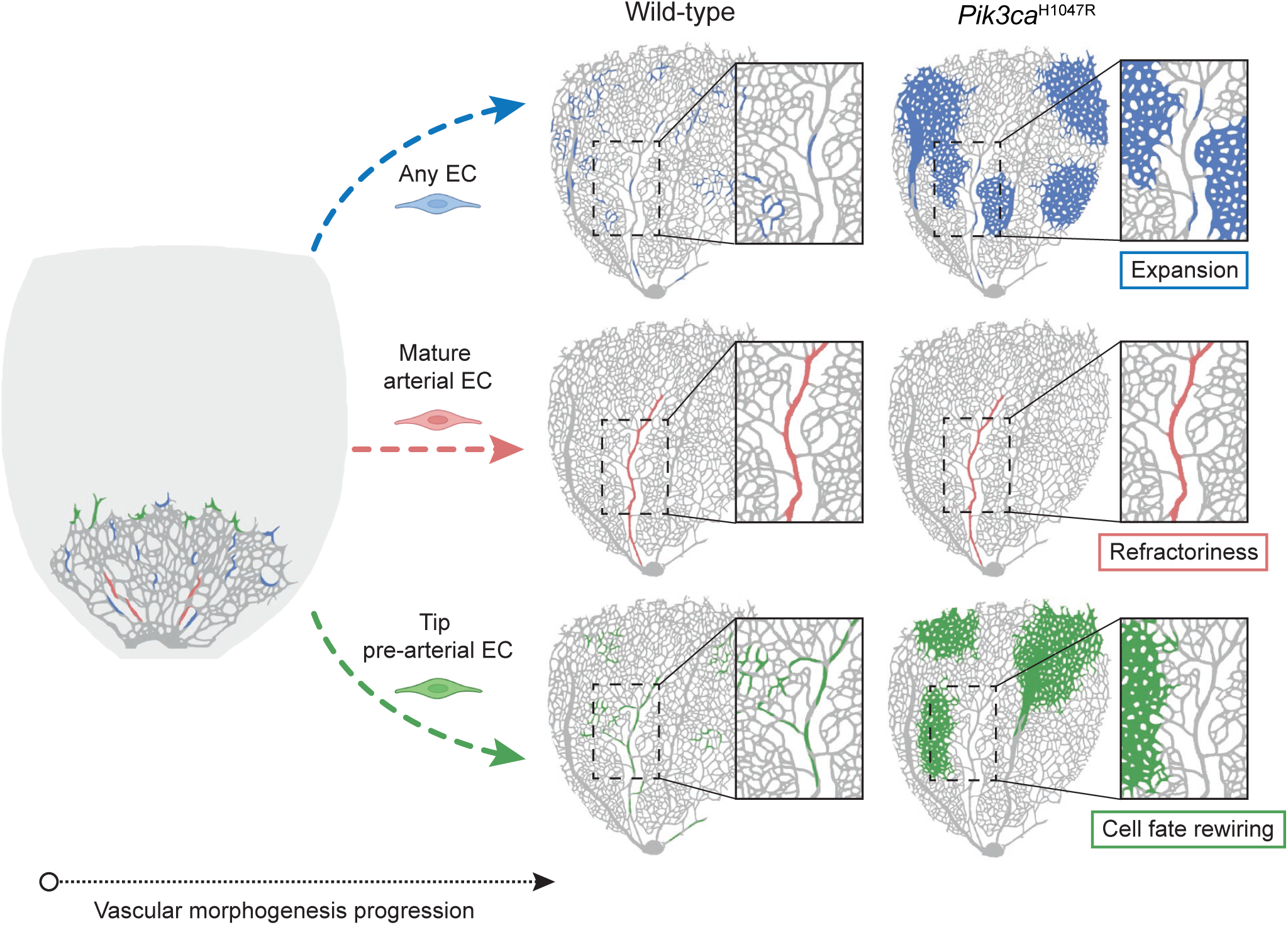
Schematic illustration of the context-dependent effects of *PIK3CA*^H1047R^ mutation in ECs. Stochastic expression of *PIK3CA*^H1047R^ in any ECs results in the generation of large clones specifically in venous and capillary ECs, causing vascular malformations while arterial ECs remain unperturbed. Expression of *PIK3CA*^H1047R^ in mature arterial ECs does not cause any pathological response. Instead, the expression of *PIK3CA*^H1047R^ in pre-arterial tip cells interrupts arterial differentiation, and induces fate switch towards venous identity. The arterial-to-venous fate switch induced by *PIK3CA*^H1047R^ is orchestrated through the upregulation of *Nr2f2*/COUP-TFII expression, a major venous fate determinant.

By combining genetic approaches that faithfully trace *PIK3CA* mutant ECs, our data demonstrate that expression of *PIK3CA* mutations is compatible with their existence in mature arteries (*Bmx* traced), albeit they do not induce any response. This is surprising, given that PI3Kα is the most active class IA PI3K isoform in ECs (Graupera et al. 2008) and is ubiquitously expressed (Vanhaesebroeck et al. 2010). Yet, we show that PI3K signaling is relatively low in arteries and is not activated in response to *PIK3CA^H1047R^*. In addition, we demonstrate that not even forced overexpression of *PIK3CA^H1047R^*can break arterial refractoriness in mature arterial cells. Building on these observations, we speculate that mature arterial ECs display cell-autonomous mechanisms to silence PI3Kα expression. Notably, the lack of arterial response to different mutations is not unique to *PIK3CA,* as other mutations, such as the loss-of-function mutation in PDCD10/CCM3 provoke similar impassive responses (Orsenigo et al. 2020), thus suggesting that arteries are also equipped with broad protective mechanisms against genetic insult.

Despite mature arterial ECs not responding to *PIK3CA^H1047R^*, this mutation interrupts the differentiation of pre-arterial cells (*i.e.,* tip cells). As soon as they emerge, tip cells are marked as pre-committed arterial ECs by expressing key arterial markers such as *CXCR4, DATCH1, HEY1, DLL4, and SOX17* (Pitulescu et al. 2017; Stewen et al. 2024; Park et al. 2021). Later, these cells migrate backward against the flow from the sprouting front to the inner plexus, settling in the artery and contributing to the growth of this vessel type. In this transition, a reduction of cell-cycle activity is required (Luo et al. 2020). We showed that upon expression of *Pik3ca^H1047R^*, tip cells are rewired to express high levels of cell cycle genes and block arterial identity. Consequently, these cells are retained in the venocapillary bed, overexpanding and forming lesions. These results align with PI3K signaling promoting venous differentiation during vascular development (Chu et al. 2016) while blocking arterial differentiation (Ang et al. 2022) and favoring stemness states (Madsen et al. 2019). Indeed, in the aging human esophagus, *PIK3CA^H1047R^* mutations in progenitor cells favor progenitor cell fate, whereas wild-type cells exhibit equal numbers of progenitor and differentiated cells per average division (Herms et al. 2024). Likewise, *PIK3CA^H1047R^* mutations evoke cell dedifferentiation during tumorigenesis in the mammary gland (Koren et al. 2015; Van Keymeulen et al. 2015).

As new mechanism findings, we discovered that *PIK3CA* mutant tip cells circumvent arterial differentiation by upregulating COUP-TFII. These data align with discoveries showing that COUP-TFII can block pre-arterial, but not arterial, specification (Su et al. 2018) and that COUP-TFII promotes stemness under proliferative pressure (Mauri et al. 2021). Despite some *Pik3ca^H1047R^*;*Nr2f2^iΔEC/iΔEC^* tip cells regaining the ability to become arterial ECs, many others remain in the capillary bed where they keep expanding. This incomplete rescue suggests that venous and proliferative programs are uncoupled downstream of *PIK3CA* mutations. This is surprising given that COUP-TFII is a master regulator of both venous identity and cell cycle progression (You et al. 2005; Wu et al. 2016) and that COUP-TFI blocks arterial differentiation by upregulating cell cycle genes (Su et al. 2018). Recent evidence has suggested that the cell cycle state determines cell fate (Chavkin et al. 2022). For instance, an early G1 cycle state is necessary to embrace a venous identity program. In contrast, arterial specification is only compatible with being in late G1. In this regard, not all tip cells exhibit a similar cell cycle state: approximately 50% of these cells are in early G1, and the remaining 50% tend to be in late G1 (Chavkin et al. 2022). Hence, it is possible that COUP-TFII depletion selectively affects tip cells in late G1, which are those more permissive to arterial signals. Instead, tip cells in the early G1 phase, and thus prone to proliferate, may already exhibit high PI3K signaling and, in turn, be less sensitive to *Nr2f2* deletion or COUP-TFII depletion. Of note, one should bear in mind that it is also conceivable that the expression of *Pik3ca^H1047R^* and deletion of *Nr2f2* occur in an asynchronous fashion with *Pik3ca^H1047R^*, thus resulting in a much faster expression than COUP-TFII depletion. This would imply that when COUP-TFII was fully depleted, many tip cells expressing *Pik3ca^H1047R^*had already clonally amplified and undergone an irreversible fate switch.

We found that both *Nr2f2* mRNA and COUP-TFII protein expression levels are increased in *Pik3ca^H1047R^* mutant ECs. This fits with previous work showing that activation of the TIE2-PI3K axis stabilizes COUP-TFII protein levels during venous specification (Chu et al. 2016). Yet, our data suggest that overactivation of PI3K signaling primarily regulates COUP-TFII at the transcriptional level. Currently, it is not understood which TFs regulate *Nr2f2* mRNA expression. Indirect evidence has suggested that SOX7, SOX18, and R-SMADs control *Nr2f2* expression in ECs (McCracken et al. 2023; Neal et al. 2019; Swift et al. 2014), yet how this connects with PI3K signaling is uncertain. Various enhancers have also been described as regulating *Nr2f2* expression in ECs. For instance, ERG is the highest expressed ETS factor in mature ECs (Shah, Birdsey, and Randi 2016) and is a crucial cognate binding TF for *Nr2f2* (Payne, Neal, and De Val 2024). However, ERG exhibits genome-wide enhancer occupancy in ECs, which has impaired understanding of how and when it promotes arterial or venous fate identity programs. Intriguingly, PI3K signaling via AKT, has been shown to modify the ERG cistrome and promote the expression of specific cell fate genes in prostate cancer (Strittmatter, Jerde, and Hollenhorst 2021). In an independent line of research, overexpression of COUP-TFII has also been found in PI3K-driven prostate tumors, with high COUP-TFII levels cooperating with PI3K signaling to sustain cancer progression (Qin et al. 2013). Based on these similarities, it is tempting to speculate that *Pik3ca^H1047R^*regulates *Nr2f2* expression via ERG in ECs.

Overall, our findings reveal that arterial ECs have a double layer of protection against *PIK3CA*. On the one hand, we identify that arterial cells are refractory to these mutations. On the other hand, we demonstrate that when *PIK3CA* mutations are expressed at early fate stages, a fate tilt towards venous and capillaries occurs, reducing the likelihood that mutant cells reach an artery. Our model shows that cell fate and differentiation stage determine the context-dependent pathological manifestations of *PIK3CA* mutations and opens the groundwork for understanding why *PIK3CA*-related tissue overgrowth is only observed in certain tissues.

## Author contribution

M.G., A.A-U., H.S. and A.R. were the main contributors to the conception, design, acquisition, and interpretation of the data, and in writing the article. J.D., A. C-R., E.C., N.T., M. A-L., S.N., J. L., P. V., performed experiments and data analysis with inputs from S.D.C., B.V., M.A.P, S.D.V., K.D.B, and R.B. L.G., and A.M-L. performed bioinformatic analysis. S.L., and E.B. liaised with human subjects and provided access to human tissue samples and clinical input for the study.

## Declaration of interests

M.G. has a research agreement with Relay Therapeutics, Inc., A Delaware corporation having a principal place of business at 399 Binney Street Cambridge, MA 02139 United States. E.B. is co-founder of Venthera; PI of the clinical trial NCT04589650 (Novartis) and Advisor for Novartis. B.V. is a consultant for Pharming (Leiden, The Netherlands) and iOnctura (Geneva, Switzerland) and a shareholder of Open Orphan (Dublin, Ireland).

## Declaration of generative AI and AI-assisted technologies in the writing process

During the preparation of this work, the author(s) used Grammarly to proofread English grammar and ChatGPT to address doubts regarding English usage. After using this tool, the author(s) reviewed and edited the content as needed and take(s) full responsibility for the content of the publication.

## Acknowledgments

We thank members of the Endothelial Pathobiology and Microenvironment Group for helpful discussions. We thank the Microscopy Core Facility and Single Cell Unit at the IJC for their support. We thank CERCA Program/Generalitat de Catalunya and the Josep Carreras Foundation for institutional support. The research leading to these results has received funding by the Spanish Ministry of Science and Innovation MICINN (PID2020-116184RB-I00 /AEI/10.13038/501100011033) and from “la Caixa” Banking Foundation under the project code LCF/PR/HR19/52160023 (also, to E.B.). The M.G. laboratory is also supported by PTEN RESEARCH Foundation (IJC-21-001); la Caixa Banking Foundation (LCF/PR/HR22/52420010 and LCF/PR/HR23/52430009); by la Asociación Española Contra el Cancer (AECC)-Grupos Traslacionales (GCTRA18006CARR); by la Fundación BBVA (Ayuda Fundación BBVA a Equipos de Investigación Científica 2019); World Cancer Research (21-0159). A.A-U. was a recipient of a fellowship from the European Union’s Horizon 2020 Research and Innovation Programme under the Marie Skłodowska-Curie grant agreement no. 101026227. L.G. was funded by the Swedish Research Council (2018-06591). H.S. is supported by the grant PRE2018-084283-MCIN/AEI and FSE, and J.D. is supported by the grant PRE2021-099260-MCIN/AEI/10.13039/501100011033 and FSE. S.D.C. was funded by la Caixa Banking Foundation Junior Leader project (LCF/BQ/PR20/11770002). R.B. was supported by the European Research Council (ERC) Consolidator Grant AngioUnrestUHD (101001814), the Ministerio de Ciencia e Innovación (PID2020-120252RB-I00), and “la Caixa” Banking Foundation (HR19-00120). M.D-L. was supported by a PhD fellowship from CNIC Severo Ochoa program (CX_E-2015-01). The CNIC is supported by the Instituto de Salud Carlos III (ISCIII), the Ministerio de Ciencia e Innovación (MCIN), and the Pro CNIC Foundation and is a Severo Ochoa Center of Excellence (grant CEX2020-001041-S funded by MICIN/AEI/10.13039/501100011033). M.A. acknowledges funding from the Wellcome Trust and The Royal Society (105942/Z/14/A). E.B. is funded by the Agencia Estatal de Investigación (Proyectos de investigación en salud PI20/00102). The authors thank the Xarxa de Bancs de Tumors de Catalunya (XBTC; sponsored by Pla Director d’Oncologia de Catalunya). We are grateful to the Band of Parents at Hospital Sant Joan de Déu for supporting the overall research activities of the Developmental Tumor laboratory (Pediatric Cancer Center Barcelona).

## Material and methods

### Mice

The *in vivo* experiments were performed in agreement with the guidelines and legislations of the Catalan Ministry of Agriculture, Livestock, Fisheries and Food (Catalonia, Spain), following protocols approved by the local Ethics Committees of the IGTP CEEAs. Mice were kept in individually ventilated cages under specific pathogen-free conditions. All mice were crossed onto the C57BL/6J genetic background.

The following mouse strains were used in this study: *Pdgfb-CreER^T2^-EGFP* (pan-endothelial Cre line) (Claxton et al. 2008); *Cdh5(PAC)CreER^T2^* (pan-endothelial Cre line) (Sörensen, Adams, and Gossler 2009), *Bmx(PAC)CreER^T2^* (arterial-endothelial Cre line) (Ehling et al. 2013) and *Esm1(BAC)CreER^T2^*(tip-endothelial Cre line) (Rocha et al. 2014), all obtained from Ralf Adams; *R26-mTmG* (Muzumdar et al. 2007) that express membrane Tomato in the absence of Cre activity and switch to membrane GFP expression upon Cre activation, allowing for the visualization and tracking of specific cell lineages; the endogenous *Pik3ca^tm1.1Waph/+^*mice, carry a germline Cre-inducible point mutation (H1047R) in one of its endogenous *Pik3ca* alleles (Kinross et al. 2012). LoxP sites flank exon 20 of the *Pik3ca* gene, and upon Cre recombination, this exon is replaced with a downstream version of exon 20 that contains a CAT to AGG alteration at codon _1047; the newly developed *Rosa26*_*CAG-LSL-H2B-Cherry/H2B-GFP-2A-HAHuPIK3CAH1047R* _mice (further_ referred to as *iFluMosaic-PIK3CA^H1047R^*), which overexpressed the human *PIK3CA^H1047R^* cDNA together with the expression of a H2B-EGFP-V5 (details in next section and Figure S4); and the *Nr2f2^flox/flox^* mouse line (Takamoto et al. 2005) kindly provided by Sophia Y. Tsai, that was used to delete *Nr2f2*.

To induce CreER^T2^ activity in pups, 4-OHT (Sigma, H7905) was dissolved in ethanol to prepare a stock solution at a concentration of 10 mg/mL. The solution was aliquoted and stored at-20°C. Pups received intragastric injections of the corresponding volume, adjusted to deliver the doses specified in the figures. To induce mosaic *Pik3ca*^H1047R^ expression in the postnatal endothelium, *Pik3ca^H1047R^*^/+^ mice were mated with *Pdgfb-CreERT2^T/+^ R26-mTmG^T/T^* mice. Pups were injected once at P1 with 0.5 mg/kg or 0.05 mg/kg 4-OHT and analysed at P4 or P6 respectively. To specifically target mature arteries, mice carrying the alleles *Pik3ca^H1047R^*^/+^, *Bmx-CreERT2^T^*^/+^ and *R26-mTmG^T/T^* were mated. The offspring were injected with 10 mg/kg of 4-OHT once at P1, P4, or P7, and subsequently analyzed at P4, P9, P12, or P21, depending on the experimental setup, as detailed in the text and figure legends. Additionally, mice carrying the *iFluMosaic-PIK3CA^T/T^* and *Bmx-CreERT2^T/+^* alleles were interbred, and the pups were injected daily from P7 to P9 with a higher dose of 4-OHT (50 mg/kg) due to the reduced recombination efficiency of this allele. To achieve tip EC-specific *Pik3ca* overactivation, mice carrying the alleles *Pik3ca^H1047R^*^/+^, *Esm1-CreERT2^T/+^,* and *R26-mTmG^T/T^* were interbred. Pups were injected with 10 mg/kg of 4-OHT at P1, and retinal analysis was performed at P2, P6, P8, or P21. Mice carrying the *R26-iFluMosaic-PIK3CA^T/T^* and *Esm1-CreERT2^T/+^*alleles were mated, and the offspring were injected with a higher dose of 4-OHT (50 mg/kg) at P1 and P2 due to the reduced recombination efficiency of this allele. For *Nr2f2* deletion specifically in tip ECs to perform rescue experiments, *Nr2f2*^flox/flox^ mice were interbred with *Pik3ca^H1047R^*^/+^, *Esm1-CreERT2^T/+^* and *R26*-*mTmG^T/T^*. Pups were injected with 25 mg/kg of 4-OHT at P1 and P2, followed by analysis of the vasculature at P6. Also, mice carrying *Nr2f2*^flox/flox^ allele were mated with *iFluMosaic-PIK3CA^T/T^*and *Esm1-CreERT2^T/+^*. In this case, mutant offspring were compared with the one resulted from the breeding of *iFluMosaic-PIK3CA^T/T^*and *Esm1-CreERT2^T/+^* mice alone. In both cases, progeny was injected with 50 mg/kg of 4-OHT at P1 and retina analysis was done at P8. To evaluate COUP-TFII protein expression in the mouse retina, offspring resulting from the cross between *Nr2f2*^flox/flox^, *Pik3ca^H1047R^*^/+^ and *Esm1-CreERT2^T/+^* mice were injected with 25 mg/kg of 4-OHT at P1 and P2, and retinal analysis was performed at P6. Cre-negative mice served as the control group. COUP-TFII staining worked optimally with a secondary antibody conjugated to a fluorophore excited in the 488 nm range, so the *R26-mTmG* alelle was excluded from this cross due to fluorescence compatibility issues. All mouse lines and primer sequences required for genotyping are provided in **Table S1.**

### Generation of the new *iFluMosaic-PIK3CA^H1047R^* mouse line

To generate the *iFluMosaic-PIK3CA^H1047R^ (Rosa26^CAG-LSL-H2B-Cherry/H2B-GFP-2A-HAHuPIK3CAH1047R^)* mouse line, we introduced the DNA insert containing the *H2B-EGFP-V5-2A-HA-humanPIK3CA^H1047R^*gene, by CRISPR/Cas9-mediated homologous-dependent repair (HDR), in the previously targeted *iChr2-Mosaic* mouse embryonic stem (ES) cell line (Pontes-Quero et al. 2017). To generate the donor construct of interest, we first obtained the *HA-huPIK3CA^H1047R^* sequence from the plasmid pBabe-puro-HA-huPIK3CA*^H1047R^* (Addgene #12524) and cloned it into plasmid SO107 (Addgene #99617)(Pontes-Quero et al. 2017), which contained the *LoxP2-H2B-EGFP-V5-2A* cassette. We then inserted the *H2B-EGFP-V5-2A-HA-huPIK3CA^H1047R^*cassette into the donor vector JP6 (Addgene #99627), which contains the hygromycin resistance gene. This resulted in the generation of the donor vector NA453, containing the complete donor construct of interest (**Extended Data Fig. 4**). The ES cells have the G4 background (George et al. 2007) and were cultured in standard ES cell media consisting of DMEM supplemented with Glutamax (31966-047, Gibco), 15% fetal bovine serum (FBS, tested for germline transmission), 1x non-essential amino acids (NEAA, Hyclone, SH3023801), 0.1% ß-mercaptoethanol (Sigma, M7522), 1x penicillin-streptomycin (Pen/Strep, Lonza, DE17-602E), and leukemia inhibitory factor (LIF). The ES cells were maintained in dishes coated with a feeder layer of mouse embryonic fibroblasts (MEFs). For each nucleofection we resuspended 2.5 million ES cells in 100 µl volume containing 1.5 µg of circular px330 plasmid (NA505) including the guide RNA sequence (Guide3, GTTGCCTATGAGAGGCTAGAC) and 3.5 µg of the circular donor plasmid (NA453) containing the *H2B-EGFP-V5-2A-HA-huPIK3CA^H1047R^*sequence and the *PGK-Hygromycin* resistance cassette (**Extended Data Fig. 4**). After nucleofection we plated 5 µl or 30 µl of the mix in two different wells on a 6-well plate with MEFs. 7 days after hygromycin selection, 24 isolated ES cell colonies were picked for storage and further screening. PCR with the flanking primers allowed us to identify ES cell clones with precise homologous recombination and insertion. After the identification of clones with precise gene targeting, we further validated the functionality by transfecting the ES cells with a Cre-expressing plasmid. Positive ES cell clones were expanded and subsequently used for microinjection into host blastocysts from the C57Bl/6J strain. Chimeric mice displaying a high percentage of agouti coat color were then bred to achieve germline transmission of the targeted insertion.

### Mouse retina isolation and whole-mount immunostaining

Mice were sacrificed by decapitation, and their eyes were carefully isolated, and incubated on ice in 4% paraformaldehyde (PFA; Sigma, 158127) in PBS for 1 h. Retinas were then dissected and fixed in 4% PFA for an additional hour on ice. After three washes with PBS, retinas were incubated overnight at 4°C in blocking buffer (1% BSA, 0.3% Triton X-100 in PBS). For COUP-TFII staining, retinas were boiled for 15 minutes in 0.01 M citrate buffer (pH 6, Sigma-Aldrich, W305600) before permeabilization. Retinas were then incubated overnight at 4°C with specific primary antibodies diluted in blocking buffer. Primary antibodies used included: anti-GFP (Abcam, ab13970; 1:100), anti-GFP (Origene, R1091P; 1:500), anti-GFP AlexaFluor 488 (ThermoFisher Scientific, A-21311; 1:200), anti-RFP CF594 (BIOTIUM, 20422; 1:200), anti-p-S6 Ser235/236 (Cell Signaling, 4857; 1:100), anti-Ki67 (Invitrogen, 14-5698-82; 1:200), anti-αSMA Cy3 (Sigma-Aldrich, C6198; 1:200), anti-Erg (Abcam, AB92513; 1:100), and anti-COUP-TFII (R&D Systems, PP-H7147-00; 1:100). After washing 3 times with 0.1% Tween 20 in PBS (PBST), retinas were incubated for 30 minutes at room temperature in PBLEC buffer [1% Triton X-100, 1 mM CaCl2, 1 mM MgCl2, and 1 mM MnCl2 in PBS (pH 6.8)]. The following secondary AlexaFluor-conjugated antibodies (1:300, ThermoFisher Scientific: A11001, A11011, A11039, A21094, A21039, A31573, S32351) were added to the retinas in PBLEC for 2 h at room temperature. Blood vessels were visualized using AlexaFluor-conjugated Isolectin GS-B4 (IB4, ThermoFisher Scientific, I21412, I32450) incubated together with the secondary antibodies. After three additional washes with PBST, retinas were flat-mounted on microscope slides with Mowiol (Calbiochem, 475904), combined with anti-fade reagent DABCO (1.4-Diazabicyclo-(2.2.2)octane; Sigma, D27802), or Fluoromount-G (SouthernBiotech, 0100-01).

### Isolation and culture of mouse lung endothelial cells

Mouse endothelial cells were isolated from the lungs of adult mice (both females and males) aged 3 to 6 weeks. Under sterile conditions in a cell culture hood, lungs were minced with a scalpel and digested in 4 U/ml dispase II (Roche, 04942078001) in Hanks’ Balanced Salt Solution (HBSS; HBSS GIBCO14170-112) for 1 h with constant agitation at 37°C. The digested tissue was dissociated by pipetting to create a single-cell suspension, and the enzyme was inactivated with Dulbecco’s Modified Eagle’s Medium (DMEM; Corning, 10-013-CV) supplemented with 10% fetal bovine serum (FBS; South America, S1810-500) and 1% penicillin-streptomycin (GIBCO, 15140-122). Cells were resuspended in PBS and incubated with rat anti-CD144/Cdh5 (BD Biosciences, 55289) antibody-coated magnetic beads for 30 minutes at room temperature. CD144-positive cells were washed with PBS containing 0.5% BSA and sorted using a magnetic separator. The sorted cells were resuspended and cultured in 12-well plates coated with 0.5% gelatin (Sigma, G1890). Cells were maintained in F12/DMEM medium (PromoCell, #C30140) supplemented with 20% fetal bovine serum (FBS), 1% penicillin-streptomycin, and 4 mL endothelial cell growth supplement/heparin (ECGF/H; Promocell C-30120) until they reached 80–90% confluency. A second selection with CD144 antibody-coated magnetic beads was performed for 1 h at room temperature. Cells were trypsinized, magnet-sorted, resuspended in F12 complete medium, and cultured further in a 6-well format. Cells were maintained at 37°C in a 5% CO2 atmosphere and used for experiments up to passage 5. Primary cells were not tested for mycoplasma contamination. To induce CreER^T2^-mediated recombination *in vitro* for Western blot or qPCR studies, ECs were treated with 2 μM 4-OHT for 6 h, while ethanol-treated cells served as controls. Recombined cells were re-seeded for experiments and cultured at 37°C in a 5% CO2 atmosphere.

### Immunofluorescence of cultured endothelial cells

Mouse lung ECs isolated from *Cdh5(PAC)CreER^T2^*; *iFluMosaic-PIK3CA^H1047R^* mice were expanded until passage 3. ECs were treated with 5 μM of 4-OHT or ethanol (control) for 6 h, followed by a 72 h incubation. A day prior to the assay, cells were split and seeded at low confluency onto gelatin-coated coverslips in 12-well plates. Four hours before collecting samples at each time point, the media was switched to starvation media. Then, cells were washed with warm 1% PBS containing Mg²⁺ and Ca²⁺ (GIBCO, 14080-055), fixed for 15 minutes in 4% PFA, and washed again with warm PBS. For staining, cells were permeabilized for 5 minutes with 0.4% Triton X-100 in PBS, followed by a 1 h incubation in blocking buffer (2% BSA in PBS). After washing with PBS, cells were incubated for 1 h at room temperature with primary antibodies diluted in blocking buffer. The primary antibodies used were anti-Ki67 (Invitrogen, 14-5698-82; 1:100), anti-p-S6 Ser235/236 (Cell Signaling, 4857; 1:100), anti-GFP (Abcam, ab13970; 1:100), and anti-RFP CF594 (BIOTIUM, 20422; 1:200). Subsequently, secondary AlexaFluor-conjugated antibodies were diluted 1:300 in blocking buffer and added to the cells for a 1 h incubation at room temperature (ThermoFisher Scientific: A11039, A21094, A21039). After additional washes with PBS, cells were incubated with DAPI (Invitrogen, D1305; 1:500) for 1 minute and then mounted on microscope slides with Fluoromount-G (Southern Biotech, 0100-01).

### Isolation, culture and sequencing of ECs from patient-derived vascular malformations

Human ECs were isolated from patient biopsies of vascular malformations. These biopsies were obtained during therapeutic surgical resection, with informed consent and approval from the Biomedical Committees at Hospital Sant Joan de Deu, Hospital Santa Creu i Sant Pau, and Hospital Universitari de Bellvitge (codes PR264/16 and PIC-96-16). Experiments adhered to the WMA Declaration of Helsinki and the Belmont Report. Data were stored in a secure database maintained by Hospital Sant Joan de Deu. Biopsies were homogenized and digested in dispase II (Roche, 04942078001) and collagenase A (Roche Diagnostics, 10103586001) for up to 1.5 h at 37°C, with continuous movement. The tissue was dissociated by pipetting into a single-cell solution, followed by enzyme inactivation with DMEM (10% FBS, 1% penicillin-streptomycin). Cells were resuspended in 0.5% BSA in PBS and incubated with CD31 antibody (Agilent Dako, M0823, clone JC70A)-coated magnetic beads (ThermoFisher Scientific, 11041) for 1 h at room temperature. The CD31+ fraction was magnetically sorted, resuspended, and cultured in 0.5% gelatin-coated wells in EGM2 medium (PromoCell, C30140) supplemented with 10% FBS, 1% penicillin-streptomycin (EGM2 complete) at 37°C and 5% CO₂ until confluency. Cells were then subjected to a second selection. Mycoplasma testing was not performed on primary cells.

To validate the presence of the mutation in the isolated ECs, genomic DNA was isolated according to the manufacturer’s protocol (ThermoFisher Scientific, K182001) and detected by NGS. For the vascular malformation (VM) #64 we obtained from 1572 reads, c.3140 A>G (H1047R) with a 45.86% variant allelic frequency (VAF) and for the VM90 from 461 reads, c.2740 A>G (E545G) with 48.37% VAF.

### Protein extraction, immunoprecipitation and immunoblotting

Cells were lysed in ice-cold lysis buffer containing 50 mM Tris-HCl (pH 7.4), 5 mM EDTA, 150 mM NaCl, and 1% Triton X-100, supplemented with protease (Roche, 11836153001) and phosphatase (Sigma-Aldrich, 4906837001) inhibitors. The protein concentration of the supernatants was measured using the Pierce BCA Protein Assay Kit (ThermoFisher Scientific, 23225) following the manufacturer’s instructions. Total cell lysates were resolved on 10% SDS-polyacrylamide gels and then transferred onto nitrocellulose membranes. The membranes were blocked with 5% milk in TBST (TBS buffer with 0.1% Tween 20) and incubated with appropriate primary antibodies diluted in 2% BSA in TBST. The following primary antibodies were used: anti-p-AKT Ser473 (Cell Signaling Technology, 4060; 1:1,000), anti-AKT (Cell Signaling Technology, 9272; 1:2,000), anti-COUP-TFII (R&D Systems, PP-H7147-00; 1:500), anti-VE-cadherin (Santa Cruz Biotechnology, sc-6458; 1:500), and anti-Vinculin (Sigma-Aldrich, 9131; 1:10,000). Secondary antibodies from DAKO were diluted in 5% milk in TBST (all at 1:5,000): swine anti-rabbit (P0399), rabbit anti-mouse (P0260), rabbit anti-goat (P0449), and rabbit anti-rat (P0450).

### cDNA synthesis and quantitative polymerase chain reaction

RNA was extracted using the Maxwell RSC SimplyRNA Cells Kit (Promega, AS1390) following the manufacturer’s instructions. The quality and quantity of the extracted RNA were assessed using a NanoDrop spectrometer. Reverse transcription was performed using 500 ng of RNA from cell lysate samples with the High-Capacity cDNA Reverse Transcription Kit (Applied Biosystems, 4368814). For quantitative polymerase chain reaction (qPCR), the QuantStudio 7 Flex system was utilized with SYBR Green real-time PCR master mix. The following primers were used for amplification: for *Nr2f2* in mouse, the forward primer was 5′-GCAAGTGGAGAAGCTCAAGG-3’ and the reverse primer was 5′-TTCCAAAGCACACTGGGACT-3’. For *NR2F2* in human samples, the forward primer was 5′-TAGTCCTGTTCACCTCAGATGCC-3’ and the reverse primer was 5′-CAGTTAAAACTGCTGCCGGAC-3’. To normalize gene expression, the following genes were used in mouse and human samples, respectively. The *ml32* gene was amplified using the forward primer 5′-ACCCCAGAGGCATTGACAAC-3’ and the reverse primer 5′-ATTGTGGACCAGGAACTTGC-3’, while *RPLP0* was amplified with the forward primer 5′-AGCCCAGAACACTGGTCTC-3’ and the reverse primer 5′-ACTCAGGATTTCAATGGTGCC-3’.

### RNA sequencing and analysis

The RNA sequencing data from *Kobialka et al.* (Kobialka et al. 2022) with public accession number PRJNA780473 (BioProject, NCBI) was reanalyzed. The counts matrix was retrieved and normalization performed using DEseq2 R package (version 1.40.2, (Love, Huber, and Anders 2014)). One biological replicate (ethanol and 4-OHT treated samples) was removed for downstream analysis to decrease variability and noise. Differential gene expression was also analyzed using DESeq2, with 0.05 as the significance cut-off, and the obtained P-values were corrected for multiple testing using the Benjamini and Hochberg method. The heatmap of selected genes was generated using ComplexHeatmaps (version 2.16.0 (Gu, Eils, and Schlesner 2016; Gu 2022)).

### Imaging analysis and quantification

Imaging was performed with Leica Stellaris 8, Leica TCS SP5, and Zeiss LSM900 confocal microscopes. Adobe Photoshop, Adobe Illustrator, ImageJ, CellProfiler 4.2.6 and Imaris 10.1 softwares were used for image editing and quantification. All images shown in the figures are maximum intensity projections. Images were taken from at least three retinal areas in each retina and are representative of at least four retinas analyzed for each genotype unless otherwise stated. The identification of the retina vascular lesions and their location was carried out based on the observation of IB4 in 10X images. Veins, arteries, and capillaries were identified by morphology as previously described in *Pitulescu et al* (Pitulescu et al. 2010). Vascular malformation incidence was quantified as the percentage of capillary beds, arterial and veins vessels exhibiting overgrowth and displayed a pathological phenotype. Retina vascularity was measured using the IB4 channel by adjusting the threshold to select the IB4+, followed by quantification of the percentage of IB4+ area in the total retinal surface. The width of arterial vessels was determined by measuring the average diameter at three different points along each artery. GFP+ cell expansion was quantified by calculating the percentage of GFP+ area within the IB4+ area. For capillary analysis, images containing only capillary beds were used. For the quantification of GFP+ cells in veins and arteries, an IB4+ area threshold was applied to identify and manually select the vessel-specific regions of interest (ROI). For the quantification of aEC-GFP+ ECs, the analyzed area was defined by a fixed ROI (200 x 630 μm at P9 and 500 x 1200 μm at P12 retinas) to maintain consistent arterial length across replicates and avoid the inclusion of other vascular types. In P21 retinas, aEC-GFP+ cells were spread across various vessel types, so the ROI for quantification included the entire vascular plexus. To quantify vascular p-S6 intensity a manual threshold was set to obtain the IB4+ area and define the ROI. Then, the integrated density of p-S6 was measured within IB4+ area. In the case of the *iFluMosaic-PIK3CA^H1047R^*, p-S6 intensity within wild-type or *PIK3CA^H1047R^* populated areas was measured by manually determining IB4+ areas containing each recombinant cell type (wild-type or *PIK3CA^H1047R^* cell clusters) and measuring the integrated density of p-S6 within ROI. In both cases, the background measurements (mean grey values) were taken from areas near the vasculature but negative for IB4. The corrected total fluorescence (CTF) was calculated using the following equation: CTF = integrated density – (vascular area x mean grey background value). EC proliferation was quantified by calculating the percentage of EC nuclei co-immunostained for both Ki67 and ERG, relative to the total number of ERG+ nuclei. COUP-TFII nuclear mean intensity was quantified through co-immunostaining for COUP-TFII and ERG. The ERG+ area was designated as the ROI, where the mean intensity of COUP-TFII was specifically measured. This mean intensity was then normalized to ERG intensity.

Quantification of Cherry-Wild-type and EGFP-*PIK3CA^H1047R^*ECs in *iFluMosaic-PIK3CA^H1047R^* retinas was performed manually, differentiating their presence in arteries, veins, and capillaries. Lesions were classified based on size as small (<10 *PIK3CA^H1047R^* cells), medium (between 10 to 35 *PIK3CA^H1047R^* cells), or large (>35 *PIK3CA^H1047R^* cells), through the identification and manual counting of mutant cells. The incidence of arteries populated with *PIK3CA^H1047R^* cells was determined as a percentage in relation to the total number of arteries containing any recombinant cell. Quantification of EC proliferation within Cherry-Wild-type and EGFP-*PIK3CA^H1047R^* cells was achieved by assessing the rate of EC nuclei immunostained for both Ki67 and either Cherry or EGFP, relative to the total number of Cherry or EGFP, respectively. Arterial vessel width in *iFluMosaic-PIK3CA^H1047R^*retinas was assessed by comparing the diameter measurement of a central area containing recombinant cells with the diameter measurement within the same vessel in a surrounding region without any recombined cell. In cultured endothelial cells, maximum intensity projections of confocal images were acquired. Image analysis was performed with CellProfiler 4.2.6. DAPI, EGFP, Cherry and Ki67 signals were used by the software as ‘primary objects’. A ‘secondary object’ dependent on DAPI was created for the quantification of pS6 (nuclei identification). CellProfiler associated pS6 signal to each nucleus and classified each cell as positive or negative for pS6. The number of wild-type and *PIK3CA^H1047R^* ECs were quantified by cross-comparison of Cherry-DAPI+ and EGFP-DAPI+ cells.

### Single-cell RNA sequencing and data processing

Mouse lung ECs isolated from *Pdgfb-CreER^T2^*-*Pik3ca^H1047R^*^/+^ mice were treated *in vitro* overnight with 2 μM 4-OHT to induce recombination and expression of *Pik3ca^H1047R^* while ethanol-treated ones were used as a control (wild-type cells for *Pik3ca*). The medium was then changed to F12 complete, and cells were collected 48 h post-induction. Cells were detached with trypsin, centrifuged, the pellet was subjected to a debris removal treatment (Miltenyi Biotec, 130-109-398), and finally cells were recollected in 0,04% BSA in PBS. For the single-cell RNA sequencing, three independent biological replicates were pooled for each treatment condition. For scRNA-seq experiments, the single cells were encapsulated in emulsion droplets using the Chromium Controller instrument using the kit 3’ Next GEM (10X Genomics). scRNA-seq libraries were prepared according to the manufacturer’s instructions. The aimed target cell recovery for each port was 10,000 cells. The generated libraries were sequenced in a NovaSeq6000 S4 (Illumina) using 2 × 150 bp paired-end read. The sequencing results for *Pik3ca^H1047R^*(4-OHT-treated) and wild-type (ethanol-treated) samples were aligned and quantified against the mouse reference genome mm10 using the Cell Ranger software (version 7.0.1) with default parameters. The output from Cell Ranger and the count matrices were read using the Read10X function from the Seurat library (version 4.9.9). Next, the established Seurat pipeline (Hao et al. 2021) was followed to analyze the samples, which were initially processed independently and later combined. As part of our data analysis, we removed genes with no reads and low-quality cells. Specifically, we excluded cells with: (i) fewer than 250 total genes, (ii) mitochondrial transcript fraction greater than 20%, and (iii) a complexity index greater than 0.8. Our quality control criteria were based on protocols from https://hbctraining.github.io/scRNA-seq/lessons/04_SC_quality_control.html and the Tonsil Atlas (Massoni-Badosa et al. 2024). To estimate doublets, which are pairs of cells sequenced under the same cellular barcode typically captured in the same droplet, we used the DoubletFinder (version 2.0.3) (McGinnis, Murrow, and Gartner 2019) package in R. This tool simulates artificial doublets using existing data and compares them with actual cells to identify potential doublets. Moreover, the DoubletFinder pipeline is integrated into the Seurat pipeline and calculates doublets per sample. After filtering, a total of 7514 cells were retained from the *Pik3ca^H1047R^* sample (4OHT-pEC-*Pik3ca*^H1047R^) and 6443 cells from the wild-type sample (ethanol-pEC-*Pik3ca*^WT^). For normalization, we applied the NormalizeData function with the LogNormalize method and a scale factor of 10,000. This process involves dividing the raw gene counts of each cell by the total counts for that cell, multiplying by the scale factor, and then performing log-normalization as log(1+x). To select the number of highly variable genes (HVG) for further analysis, we first calculated the distribution of non-zero counts across all genes using the RNA assay data. This was achieved by applying the colSums function from the Matrix package (v 1.6.0) (Bates, Maechler, and Jagan 2000). We then identified the number of HVG by adding 100 to the third quartile of this distribution, establishing a threshold for variability selection. Following normalization, we performed a z-score transformation on the normalized values using Seurat’s ScaleData with default parameters, followed by principal component analysis using RunPCA. Finally, clustering analysis was performed based on the edge weights between any two cells, utilizing a shared nearest-neighbor graph produced by the Louvain algorithm, implemented in Seurat’s FindNeighbors and FindClusters functions. Louvain clustering was performed at resolution 0.3 and clusters were annotated based on known tip, venous, arterial, and proliferative markers as described in the main text. Taking advantage of the PCA dimensionality reduction performed as part of the pre-processing pipeline, cell types were represented in PCA dimensions 1 and 2 to assess the degree of similarity between different clusters. Using this representation, we conducted a differential gene expression analysis employing the Wilcoxon test to compare *Pik3ca^H1047R^* (4OHT-pEC-*Pik3ca*^H1047R^) with the wild-type sample (ethanol-pEC-*Pik3ca*^WT^) within the Tip/Arterial-Venous cluster (C1/C2). This analysis identified the genes differentially expressed in the *Pik3ca^H1047R^* ECs. Statistically significant genes (Bonferroni-adjusted p-value < 0.05) with positive or negative log2FC values were selected as the set of significantly regulated genes. Violin plots of the selected genes were generated using the VlnPlot function from the Seurat package, with the split.by argument applied to divide the average expression of the genes. Heatmaps of gene expression were created using the ComplexHeatmap library in R (Gu, Eils, and Schlesner 2016). The heatmap values represented the scaled and normalized expression levels of the genes within the Tip/Arterial-Venous cell cluster (C1/C2).

## Code availability

No new algorithms were developed for this article. All original code has been deposited at GitHub (https://github.com/anemartinezlarrinaga2898/Sabata_Rocat_et_al.git).

## Data availability

The scRNA-seq expression data supporting is uploaded to the Gene Expression Omnibus (GEO) repository under the accession ID GSE287780. RNA-seq data was obtained from the BioProject repository (NCBI) under the accession ID PRJNA780473 (https://www.ncbi.nlm.nih.gov/bioproject/PRJNA780473) (Kobialka et al. 2022).

## Statistics

Statistical analysis was performed using Prism 10 (GraphPad Software Inc.). All data are displayed with individual data points that indicate biological replicates and presented as mean ± s.d. or mean ± s.e.m. (as indicated in the figure legends). At least four biological replicates were used. Statistical significance between two groups was assessed using either an unpaired two-tailed Student’s t-test or the nonparametric Mann-Whitney test. For comparisons involving more than two groups, one-way and two-way ANOVA followed by Tukey’s post-hoc test for multiple comparisons were used. No statistical method was employed to predetermine sample sizes; sample sizes were based on prior experiments. Reproducibility was confirmed through multiple independent experiments. The experiments were not randomized, and the investigators were not blinded during the experiments or data analysis.

## Antibodies

anti-GFP - Abcam, ab13970; 1:100 (IF)

anti-GFP - Origene, R1091P; 1:500 (IF)

anti-GFP AlexaFluor 488 - ThermoFisher Scientific A-21311; 1:200 (IF)

anti-RFP CF594 - BIOTIUM, 20422; 1:200 (IF)

anti-p-S6 Ser^235/236^ - Cell Signalling, 4857; 1:100 (IF)

anti-Ki67 - Invitrogen, 14-5698-82; 1:200 (IF)

anti-αSMA Cy3 - Sigma-Aldrich, C6198; 1:200 (IF)

anti-Erg - Abcam, AB92513; 1:100 (IF)

anti-COUP-TFII - R&D Systems, PP-H7147-00; 1:100 (IF) 1:500 (WB)

anti-p-AKT Ser^473^ - Cell Signaling Technology, 4060; 1:1,000 (WB)

anti-AKT - Cell Signaling Technology, 9272; 1:2,000 (WB)

anti-VE-cadherin - Santa Cruz Biotechnology, sc-6458; 1:500 (WB)

anti-Vinculin - Sigma-Aldrich, 9131, 1:10,000 (WB)

## Extended Data Figures

**Extended Data Fig. 1.**
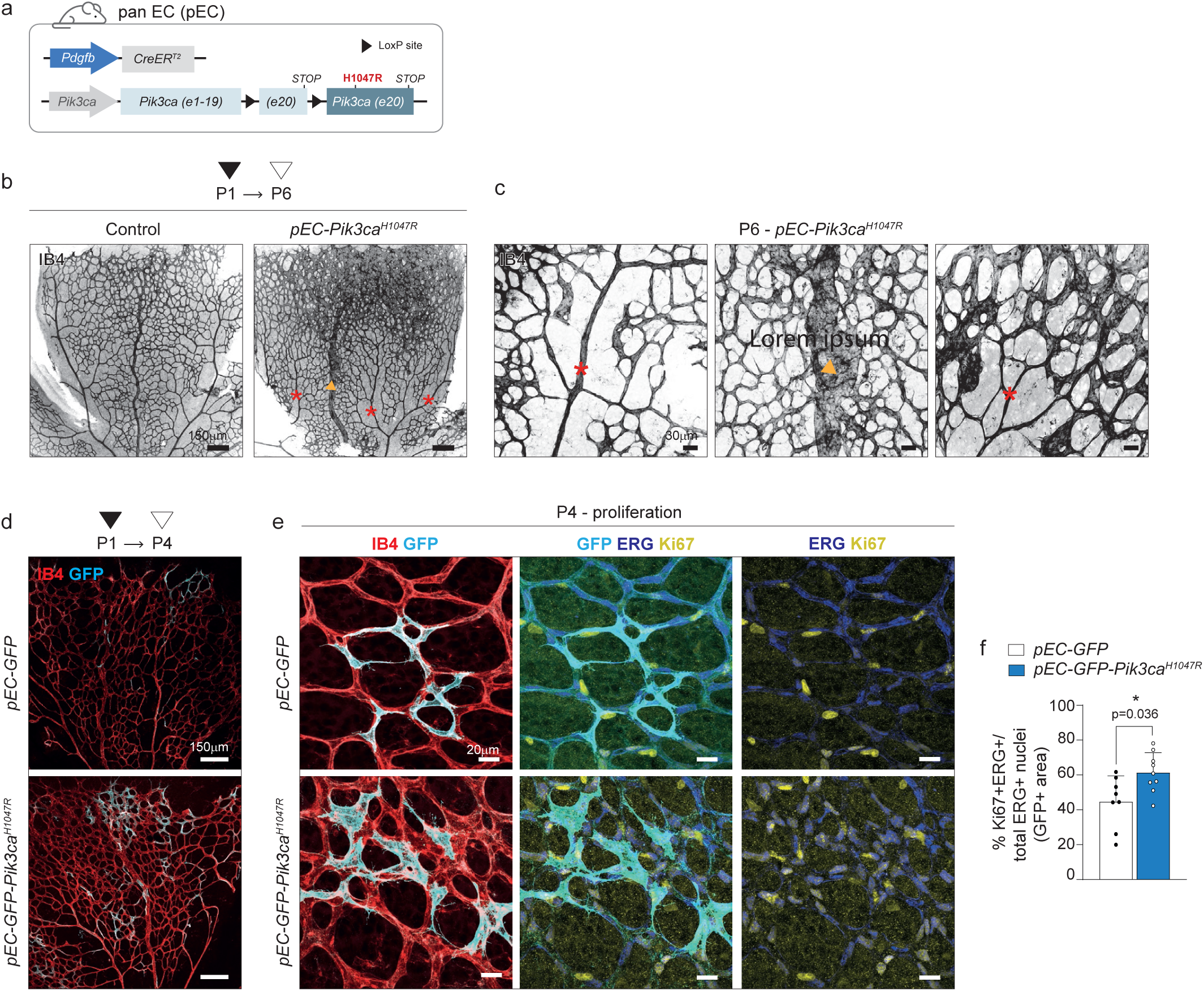
Phenotype overview and assessment of EC proliferation in *Pik3ca^H1047R^* retinas related to. Figure 1 (a) Schematic illustration of the transgenic mouse strategy combining the pan-endothelial (pEC) tamoxifen-inducible Cre (*Pdgfb-CreER^T2^*) allele with *Pik3ca^H1047R^* allele (*pEC-GFP-Pik3ca^H1047R^*). (b) Representative confocal images of control (Cre-) and *pEC-Pik3ca^H1047R^* P6 retinas. IB4 staining shows that only veins (orange arrowhead) and capillaries, but not arteries (red asterisks) develop vascular abnormalities upon *Pik3ca^H1047R^* expression. (c) High-magnification confocal images of arteries (red asterisks), vein (orange arrowhead), and capillary plexus (right panel) from P6 *pEC-Pik3ca^H1047R^* retinas stained with IB4. (d) Representative confocal images of *pEC-GFP* and *pEC-GFP-Pik3ca^H1047R^* P4 retinas stained with IB4 (red, blood vessels) and GFP (cyan, tracing recombined ECs). Retinas were isolated at P4, 3 days after 0.5 mg/kg 4-OHT injection at P1. (e) High-magnification confocal images of P4 retinas stained for IB4 (red, blood vessels), GFP (cyan, recombined cells), ERG (blue, EC nuclei), and Ki67 (yellow, proliferative cells). At this stage, *pEC-GFP-Pik3ca^H1047R^* retinas show initial vascular lesions. (f) Quantification of EC proliferation within the GFP+ area, represented as the percentage of ERG^+^ Ki67+ nuclei per total number of ERG+ nuclei (n ≥ 8 retinas per genotype). Data are presented as mean ± s.d. Statistical analysis was performed using nonparametric Mann-Whitney test. Scale bars: 150 µm (b, d), 30 µm (c) and 20 µm (e).

**Extended Data Fig. 2.**
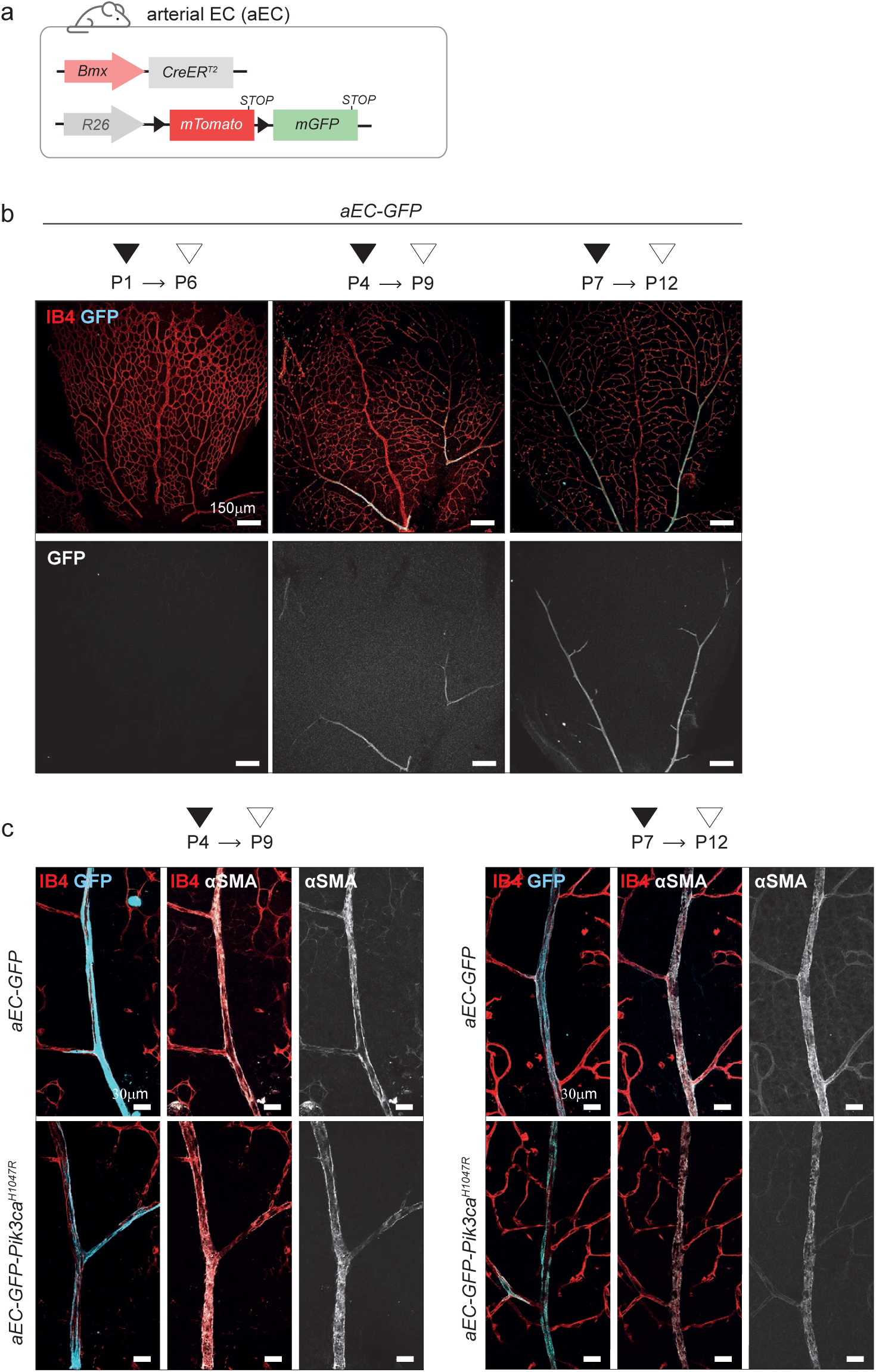
Tracing *Bmx*+ ECs at different time points and impact of *Pik3ca^H1047R^* expression in αSMA staining related to. Figure 2 (a) Schematic illustration of the transgenic mouse strategy combining the arterial EC-specific (*aEC*) tamoxifen-inducible Cre (*Bmx(PAC)*-CreER^T2^) allele with the *R26*-*mTmG* reporter allele (*aEC-GFP*). (b) Representative confocal images of *aEC-GFP* retinas after 10 mg/kg of 4-OHT treatment at P1, P4 or P7, followed by retina isolation and analysis at P6, P9 or P12, respectively. IB4 (red, blood vessels) and GFP (cyan, recombined cells) immunostaining show no GFP+ ECs when induced at P1, indicating a lack of *Bmx* expression at this stage. In contrast, GFP+ cells are observed after induction at P4 and P7. (c) High-magnification images of arteries from *aEC-GFP* and *aEC-GFP*-*Pik3ca^H1047R^* P9 (left) and P12 (right) retinas stained with IB4 (red, blood vessels), anti-GFP (cyan, recombined cells) and anti-αSMA (white, marker of smooth muscle cells). αSMA staining is used to identify arteries and assess their structural integrity. Scale bars: 150 µm (b) and 30 µm (c).

**Extended Data Fig. 3.**
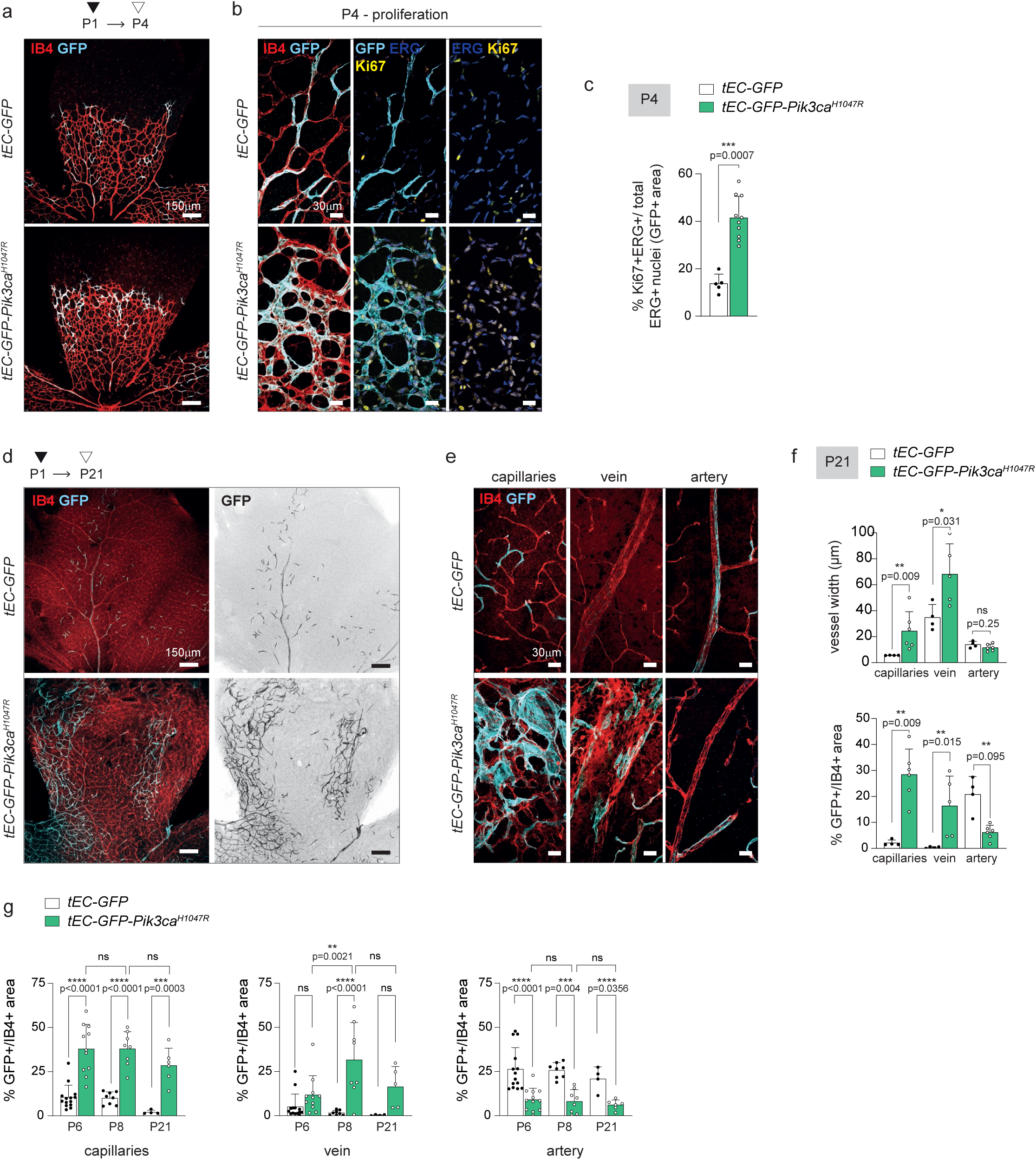
Assessment of EC proliferation and long-term clonal dynamics of tip ECs related to. Figure 3 (a) Representative confocal images of P4 *tEC-GFP* and *tEC-GFP-Pik3ca^H1047R^* retinas treated with 10 mg/kg 4-OHT at P1, stained for IB4 (red, blood vessels) and GFP (cyan, recombined cells). At P4, incipient vascular malformations are detected. (b) High-magnification confocal images of P4 retinas stained for IB4 (red, blood vessels), GFP (cyan, recombined cells), ERG (blue, EC nuclei), and Ki67 (yellow, proliferative cells). (c) Quantification of EC proliferation within the GFP+ area in *tEC-GFP* and *tEC-GFP-Pik3ca^H1047R^*P4 retinas, represented as the percentage of ERG+Ki67+ nuclei per total number of ERG+ nuclei (n ≥ 5 retinas per genotype). (d) Representative confocal images of P21 *tEC-GFP* and *tEC-GFP-Pik3ca^H1047R^* retinas treated with 10 mg/kg 4-OHT at P1, stained for IB4 (red, blood vessels) and GFP (cyan, recombined cells). At P21, the long-term behavior of P1 *Esm1*-derived progeny can be assessed. (e) High-magnification confocal images of capillaries, veins and arteries in P21 retinas stained for IB4 (red, blood vessels) and GFP (cyan, recombined cells). (f) Quantification of vessel width (top) and percentage of GFP+ area (bottom) in *tEC-GFP* and *tEC-GFP*-*Pik3ca^H1047R^* P21 retinas, specifically in capillaries, veins, and arteries (n ≥ 4 retinas per genotype). (g) Quantification of the percentage of GFP+ area in *tEC-GFP* and *tEC-GFP*-*Pik3ca^H1047R^* P6, P8 and P21 retinas, specifically in capillaries, veins, and arteries. Data previously presented in Fig. 3f and Extended Data Fig. 3f, now represented differently to illustrate temporal changes. Data are presented as mean ± s.d. Statistical analysis was performed using nonparametric Mann-Whitney test (c, f) and 2-way ANOVA with Sidak’s correction for multiple comparisons (g). ns, non-statistical. Scale bars: 150 µm (a, d) and 30 µm (b, e).

**Extended Data Fig. 4.**
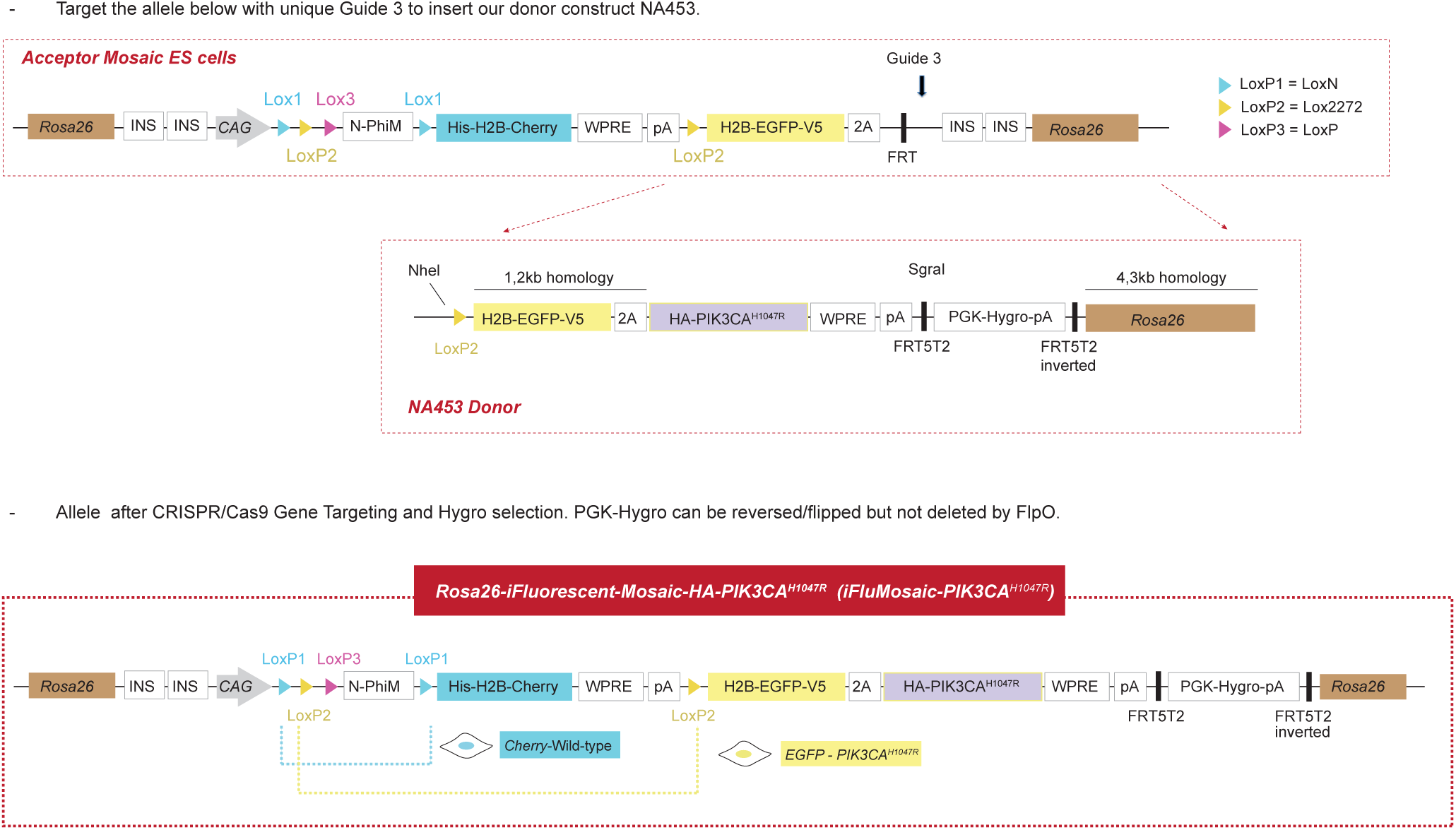
Summary of the steps performed to generate the new *Rosa26-iFluorescent-Mosaic-PIK3CA^H1047R^* mouse model, related to. Figure 4 All details in M&M. PGK-hygro-pA, resistance marker for selection; WPRE-pA, viral sequence to enhance the stability and efficiency of transgene expression by improving the export of mRNA from the nucleus to the cytoplasm; CAG, Strong and ubiquitous promoter; PGK-Neo-pA, resistance marker for ES cell selection; 2A, viral peptide allowing equimolar expression of multiple independent proteins from a single ORF; HA, V5, and His are small epitopes that can be used for specific antibody detection; H2B, histone tag that targets proteins to the chromatin/nucleus; N-PhiM, non-fluorescent protein that is used as a reporter of promoter expression; INS, Insulators, are DNA sequences that regulate gene expression by preventing enhancers from activating nearby genes.

**Extended Data Fig. 5.**
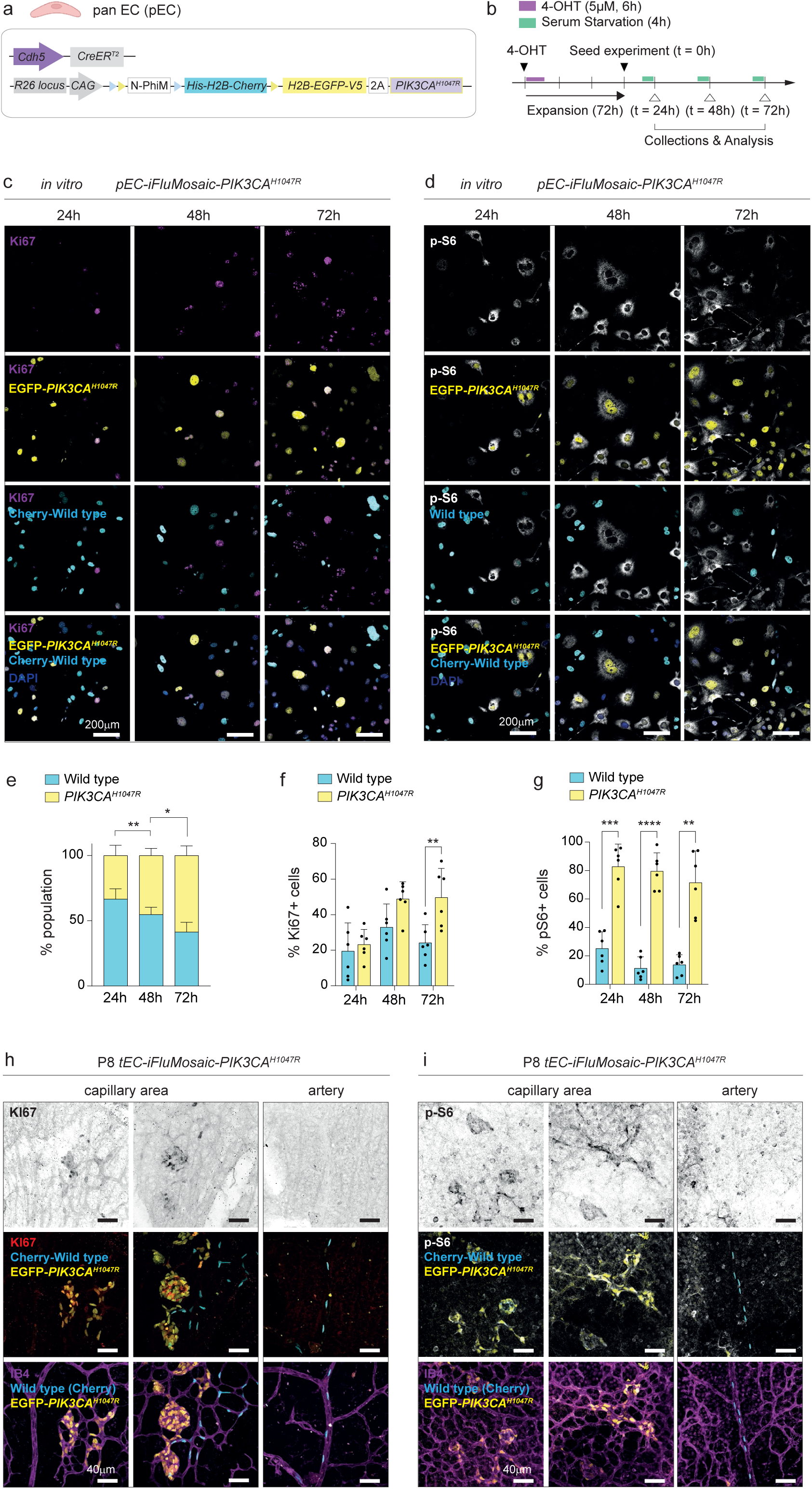
***In vitro* validation of the *iFluMosaic-PIK3CA^H1047R^* model and representative images of *iFluMosaic-PIK3CA^H1047R^* mouse retinas stained for different markers, related to** Figure 4 (a) Schematic illustration of the transgenic mouse strategy combining the pan EC-specific (pEC) tamoxifen-inducible Cre (*Cdh5(PAC)-CreER^T2^*) allele with the *iFluMosaic-PIK3CA^H1047R^* allele. (b) Strategy for *in vitro* EC expansion, 4-OHT administration and analysis. 6 h 4-OHT treatment was followed by 72h cell expansion. Daily (every 24h) collections were preceded by 4h of complete serum starvation. (c, d) Representative confocal images of primary mouse lung ECs isolated from *pEC-iFluMosaic-PIK3CA^H1047R^* mice collected at 24, 48, and 72 h after seeding. Cells were stained for DAPI (nuclei, blue), EGFP (*PIK3CA^H1047R^* expressing ECs, yellow), Cherry (wild-type ECs for *Pik3ca*, cyan), and Ki67 (proliferative cells, magenta) (c) or phospho (p)-S6 (Ser235/236) (grey) (d). (e, f, g) Quantification of Cherry-Wild-type (cyan) and EGFP-*PIK3CA^H1047R^* (yellow) recombined ECs at 24, 48, and 72 h post seeding (e). Shown as percentage of *PIK3CA^H1047R^* and wild-type cell sum. Within each population, quantification of the percentage of cells positive for Ki67 (f) or p-S6 (g) (n = 6 biological replicates). (h,i) High-magnification images of capillaries and arteries from P8 *tEC-iFluMosaic-PIK3CA^H1047R^* retinas showing immunodetection of IB4 (magenta), Cherry (cyan, wild-type ECs for *Pik3ca*), EGFP (yellow, *PIK3CA^H1047R^* expressing ECs) and Ki67 (red, proliferative cells) (h) or p-S6 (white) (i). Data are presented as mean ± s.d. 2-way ANOVA statistical analysis was performed followed by Tukey test for multiple comparisons (e, f, g). Scale bars: 200 µm (c, d) and 40 µm (h, i).

**Extended Data Fig. 6.**
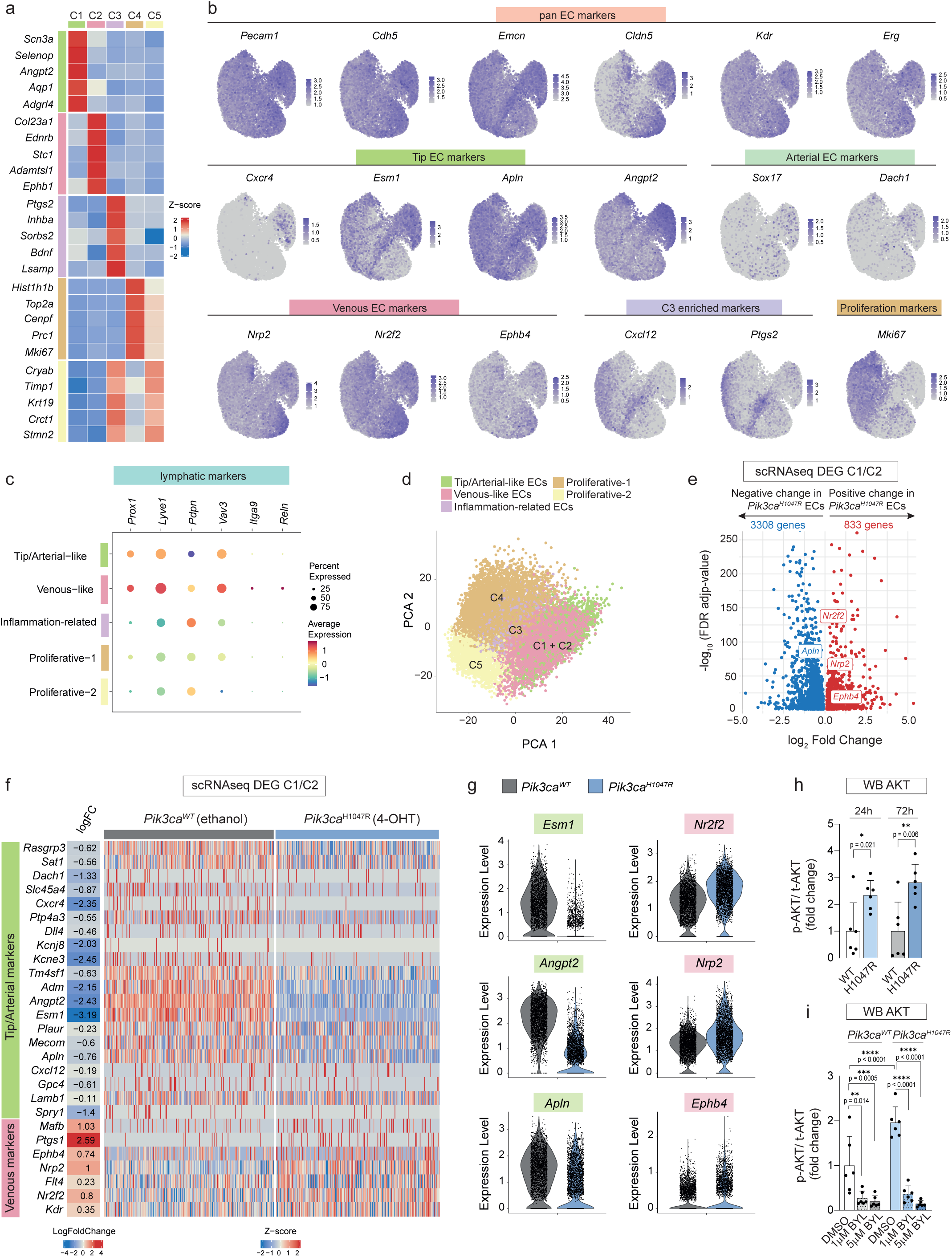
Extended analysis supporting. Figure 5 (a) Heatmap representing Z-normalized gene expression levels of the top 5 most highly expressed markers of each cluster. (b) Feature plots showing the gene expression of known endothelial-specific marker genes and key markers of each cluster. (c) Dot plot showing the average expression of lymphatic markers for each cluster. (d) PCA plot of merged *Pdgfb-CreER^T2^* (pEC)-*Pik3ca^H1047R^* mouse lung endothelial cells (ECs) 48 h after treatment with ethanol (vehicle, wild type for *Pik3ca*) and 4-OHT (expressing *Pik3ca^H1047R^*) derived from scRNA-seq data. Cells are colored based on clusters identified in the previous UMAP representation in Fig. 5a. C1 (tip-like ECs) and C2 (venous-like ECs) clusters which were found enriched in *Pik3ca^WT^* and *Pik3ca^H1047R^* ECs respectively, clustered together here (C1/ C2), allowing for the following comparisons between genotypes. (e) Volcano plot of genes analyzed by scRNAseq in C1 and C2 clusters upon *Pik3ca^H1047R^* expression. Differentially expressed genes (DEGs) (Padj < 0.05) in *Pik3ca^H1047R^* over *Pik3ca^WT^* ECs from C1 and C2 (grey, unchanged; blue, downregulated; red, upregulated). (f) Heatmap showing the gene expression of selected DEG per cells from C1/C2 cluster in *Pik3ca^WT^* and *Pik3ca^H1047R^* ECs. (g) Violoin plots of selected DEG between *Pik3ca^WT^* vs *Pik3ca^H1047R^* conditions in C1/C2 cluster. (h) Quantification of phopsho(p)-AKT (Ser473) protein levels normalized to total AKT levels (related to Figure 5G) (n = 6 biological replicates). (i) Quantification of p-AKT (Ser473) protein levels normalized to total AKT levels (related to Figure 5J) (n = 6 biological replicates). Data are presented as mean ± s.d. Statistical analysis was performed using an unpaired t-test (h) and 2-way ANOVA followed by Tukey test for multiple comparisons (i).

**Extended Data Fig. 7.**
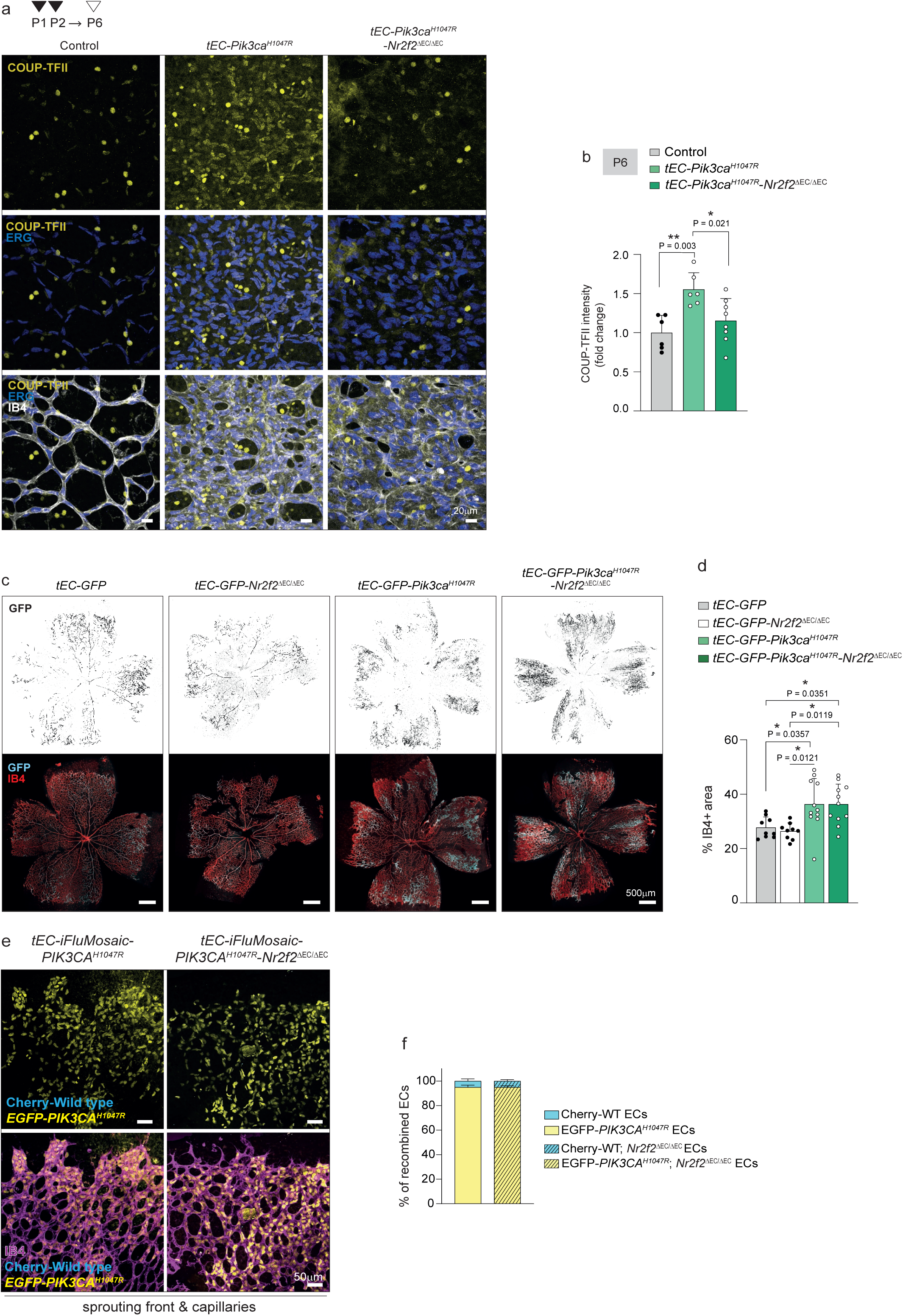
Validation of effective *Nr2f2* deletion and its impact on vascular area. (a) Representative confocal images of control, *tEC-GFP-Pik3ca^H1047R^* and *tEC-Pik3ca^H1047R^*-*Nr2f2^ΔEC/ΔEC^* P6 mouse retinas after 25 mg/kg 4-OHT induction at P1 and P2. Immunostaining for IB4 (white), ERG (blue, EC nuclei,), and COUP-TFII (yellow) showing upregulation of COUP-TFII in *Pik3ca^H1047R^* vessels. (b) Quantification of COUP-TFII intensity within ERG+ areas (EC nuclei) in capillary vessels from control, *tEC-Pik3ca^H1047R^* and tEC*-Pik3ca^H1047R^*-*Nr2f2^ΔEC/ΔEC^* P6 retinas (n ≥ 6 retinas per genotype). (c) Representative confocal images of *tEC-GFP*, *tEC-GFP-Nr2f2^ΔEC/ΔEC^*, *tEC-GFP-Pik3ca^H1047R^*, and *tEC-GFP-Pik3ca^H1047R^*-*Nr2f2^ΔEC/ΔEC^* P6 mouse retinas, stained for IB4 (red, blood vessels) and GFP (cyan, P1 and P2 *Esm1*-derived progeny). (d) Quantification of total vessel area (IB4+ area) comparing *tEC-GFP*, *tEC-GFP-Nr2f2^ΔEC/ΔEC^*, *tEC-GFP-Pik3ca^H1047R^*, and *tEC-GFP-Pik3ca^H1047R^*-*Nr2f2^ΔEC/ΔEC^* of P6 retinal vasculature (n ≥ 9 retinas per genotype). (e) Representative confocal images of *tEC-iFluMosaic-PIK3CA^H1047R^* and *tEC-iFluMosaic-PIK3CA^H1047R^-Nr2f2^ΔEC/ΔEC^* P8 retinas stained for IB4 (magenta, blood vessels), Cherry (cyan, wild-type ECs for *Pik3ca*) and EGFP (yellow, *PIK3CA^H1047R^*expressing ECs) at the capillary front. (f) Quantification of the total number of Cherry-Wild-type and EGFP-*PIK3CA^H1047R^* ECs in *tEC-iFluMosaic-PIK3CA* and *tEC-iFluMosaic-PIK3CA-Nr2f2^ΔEC/ΔEC^* P8 retinas (n = 16 retinas per genotype). Data are presented as mean ± s.d. Statistical analysis was performed using one-way ANOVA, followed by Tukey test for multiple comparisons (b, d) and 2-way ANOVA with Sidak’s correction for multiple comparisons (f). Scale bars: 20 µm (a), 500 µm (c), and 50 µm (e).

---

**Table S1.**
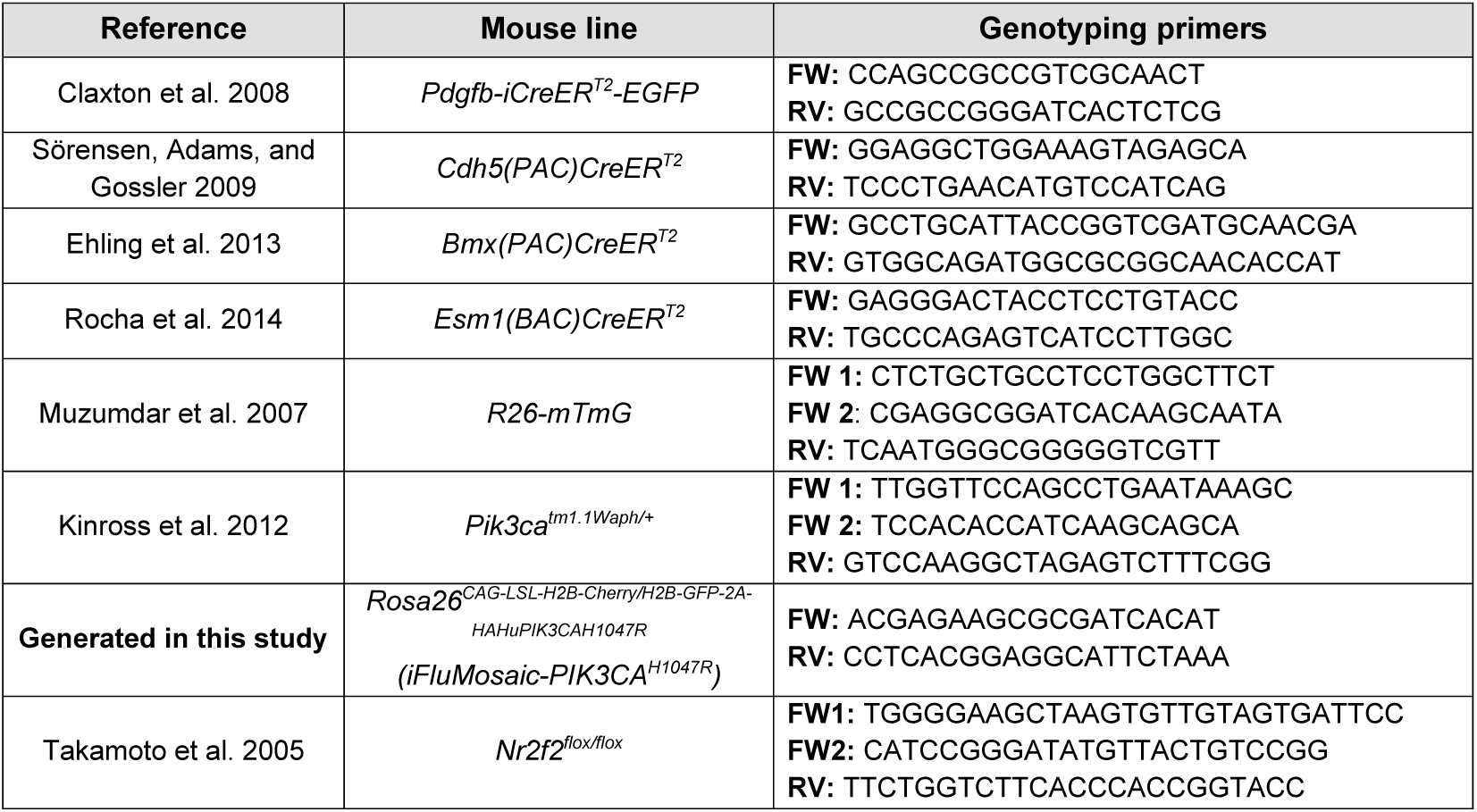
Mouse lines and genotying primers.

